# Multilayered human activities shape the microbial communities of groundwater-dependent ecosystems in an arid oceanic island

**DOI:** 10.64898/2025.12.01.691164

**Authors:** Francesco Di Nezio, Andrea Di Cesare, Marta García-Cobo, David Brankovits, Raffaella Sabatino, Giulia Borgomaneiro, Zacarías Fresno-López, Márcia Neunschwander Kurtz, Sarah Boulamail, Francesco Cozzoli, Lara Fumarola, Brett C. Gonzalez, Alvaro Roldán, Cristina Camacho, Alvaro García-Herrera, Leopoldo Moro, Concepcion Valdivia, Elena Mateo, Guillermo García-Gómez, Diego Fontaneto, Gianluca Corno, Ester M. Eckert, Alejandro Martínez

## Abstract

Island coastal aquifers, though physically small compared to continental groundwater systems, are of huge ecological and societal importance, sustaining functions that connect to locally crucial provision, maintenance and culture ecosystem services. Those functions are largely dependent on the presence of highly adapted biological communities, for which, their microbial communities remain understudied. Our goal is to describe the bacterial communities across the groundwater-dependent ecosystems on Lanzarote, spanning a gradient of anthropogenic pollution using 16SrRNA amplicon sequencing. We sampled coastal caves and pools, wells and water galleries, springs, saltworks and marine bays affected by submarine groundwater discharge. Ecological analyses highlight that richness and composition of bacterial communities strongly depend on the type of habitats. Pathogens and human-derived species were ubiquitous in our samples, but, strikingly, caves and wells were strongly enriched with them compared to other habitats. We propose that our results highlight the susceptibility of groundwater environments to pollution and indicate that aquifers act as reservoirs of biological contamination in addition to natural diversity—regardless of their salinity. Since this enrichment might compromise some of the functions and services that groundwater-dependent ecosystems provide in oceanic islands, we call for integrative conservation strategies that include hydrological and biological perspectives into the decision making.

## 1. Introduction

Islands, through time, have been both cradles and reservoirs of biodiversity (Gillespie and Clague, 2009). Although islands cover only 5.3% of Earth’s land surface, they not only harbour roughly 20% of described species, with exceptionally high endemism rates, but also 27% of human languages (Tershy et al., 2015; Fernández-Palacios et al., 2021). As much as ecosystems, human societies have flourished on islands, channeling oceanic trade and migration and inspiring literature and legends. Yet, all these natural and cultural heritage ultimately rely on insular aquifers: groundwater systems that store and circulate the scarce, drinkable freshwater that makes island’s terrestrial life possible. Because surface freshwater is often scarce and strongly dependent on precipitation, island societies have depended on aquifers for drinking water and irrigation. Island aquifers also sustain vegetation and wetlands along altitudinal gradients, buffering droughts effects, and coastal groundwater-fed pools and caves hold cultural heritage and serve as touristic attractions. Beyond the island itself, groundwater discharge into the ocean regulates coastal ecosystems by delivering nutrients and trace elements (Moore, 2010; Moosdorf et al., 2015; Luijendijk et al., 2020), and shaping coastal food webs (Gibson et al., 2022; Mammola et al., 2025).

Stored in volcanic or carbonate rocks, island aquifers are stratified systems, with a meteoric freshwater lens overlying denser, recirculating seawater. At their interface, a subterranean estuary develops, creating a biogeochemical reaction zone that modulates the chemistry of groundwater before its discharge to the coastal sea (Moore, 1999; Santos et al., 2021). On volcanic islands, these hydrological systems establish early and change through the island’s ontogeny. On young, high-altitude islands, abundant orographic rainfall sustains recharge, feeding perched aquifers on the mountain flanks that coalesce downslope into a thick basal meteoric lens above the saline aquifer. Over time, as islands erode and subside (Whittaker et al., 2008), elevation and recharge decline and parts of the aquifer become effectively disconnected from modern inputs. At this stage, groundwater in perched aquifers may become non-renewable on human time scales (i.e. fossil), while the basal meteoric lens progressively thins, density stratification weakens, seaward groundwater flux diminishes, and overall salinity increases (van Hengstum et al., 2019).

Beyond the ecosystem-level salinity gradients of a subterranean estuary, interconnected groundwater-dependent habitats associated with coastal aquifers are characterized by further local-scale gradients in salinity, oxygen, and nutrients (Jang et al., 2012; Cho et al., 2018; Kim et al., 2019; Moore and Joye, 2021). First documented in tidal pools (Holthuis, 1973), and later explored through the extensive cave networks connected to them (Stock, 1986), these ‘anchialine habitats’ were initially studied for their highly specialized, endemic fauna, including lineages with apparent deep sea affinities or interpreted as ‘living fossils’ (Iliffe and Kornicker, 2009). In these habitats, microbes have a key role in sustaining nutrient and carbon cycling, and shuttling carbon and energy to the higher trophic levels of the anchialine food web (Brankovits et al., 2017; Ruiz-González et al., 2021; Elbourne et al., 2022). In hypersaline areas, halophilic archaea and halotolerant bacteria employ compatible-solute accumulation or “salt-in” ion-balancing strategies to withstand osmotic stress (Narasingarao et al., 2012; Vaidya et al., 2018). In springs, anchialine pools, and sinkholes, phototrophs and rhodopsin-bearing prokaryotes harvest light. In the darkness of caves, bacterial communities rely on limited organic inputs from the outside or from in situ production by microbes that utilize reduced sulfur, methane, nitrogen, or other compounds (Pohlman, 2011; Kajan et al., 2022; Meland et al., 2023).

Coastal aquifer communities are highly vulnerable to environmental change. At local and regional scales, tourism, agriculture, and urbanization intensify groundwater extraction, nutrient enrichment, and contamination (Kochary et al., 2018; Dey et al., 2022; Dong et al., 2024). At broader scales, climate change and sea-level rise drive temperature increase, promote saltwater intrusion, and intensify drought (Song and Zemansky, 2012; Oberle et al., 2017; Chesnaux et al., 2021). Altogether, these impacts disrupt both faunal and microbial communities, alter trophic interactions, reduce functional diversity, and accelerate ecosystem degradation. Because bacterial communities respond rapidly to stress, they are early warning indicators of shifts in water quality and geochemical stability. Rising ocean and groundwater temperatures can increase microbial activity (Benz et al., 2024), and a large fraction of known pathogenic microbes may gain ecological advantages under future climates (Carlson et al., 2022; Mora et al., 2022). Concurrently, reduced oxygen solubility exacerbates hypoxia and anoxia, conditions that can favor anaerobic pathogens (Cuthbert et al., 2019; Fatima, 2024). In combination with nutrient enrichment and hydrological stress, warming is likely to intensify ecological degradation and elevate health risks through groundwater contamination by pathogens and fecal bacteria (He et al., 2018; Lyons et al., 2021). Those scenarios reinforce the urgency of baseline microbial datasets, robust early warning systems, and timely policy interventions (Ferguson and Gleeson, 2012). Yet, many aquifers lack a robust baseline description of their actual microbial community, limiting our ability to design research roadmaps and integrative conservation strategies.

The goal of this paper is to describe the bacterial communities across the groundwater-dependent ecosystems on Lanzarote (Canary Islands, Spain), spanning a gradient of anthropogenic pollution. We selected Lanzarote because the low-altitude coastlines expose the basal aquifer through lava tubes and anchialine pools, whereas perched aquifers are easily accessible through springs and wells. These waters have long underpinned human settlement, navigation and coastal trade, as well as supporting a uniquely diverse anchialine fauna, but accelerating population growth together with pollution, rising water demand, and urban expansion, now threatens Lanzarote’s groundwater ecosystems (Martinez et al., 2016). The main hypothesis underlying this research is that habitat differences structure bacterial communities through the combined effect of environmental and anthropogenic processes. Environmental-related bacterial taxa, which have colonized the island since its formation, should reflect both island-level and local gradients present in groundwater and related ecosystems. We expect their richness to vary with resource availability and salinity, with key drivers including light and distance from the sea, and their composition to be dominated by turnover (environmental selection). In contrast, anthropogenically derived taxa, introduced directly or indirectly via humans, livestock, crops, and materials, should display local enrichment linked to land use and hydrological connectivity. Therefore, we predict their richness to be higher in impacted habitats and their composition to be best described by nested patterns. Finally, we anticipate that more distinct habitats, ecologically or due to the presence of strong anthropogenic impacts, will present more distinct ecological communities. By involving both researchers and stakeholders in our team, we translate these results into a set of consensus-based strategies aimed at enhancing the monitoring and conservation of Lanzarote’s groundwater-dependent habitats.

## Materials and methods

### 2.1. Study sites and sampling design

To test the hypothesis that different groundwater-dependent habitats harbor distinct bacterial communities, we sampled the main accessible aquifer-related habitats of Lanzarote, including lava-tube caves, anchialine pools, saltworks, and groundwater wells and galleries. These sites represent access to both pristine and impacted aquifers, encompassing a documented history of anthropogenic use ranging from pre-Hispanic exploitation to recent tourism and agriculture. Table 2 provides an overview of the aquifer compartments, representative habitats, ecosystem functions, and associated anthropogenic impacts considered in this study.

Sampling was conducted across 25 sites representing distinct habitat types on Lanzarote (Figure 2, Table 1).

**Figure 1.**
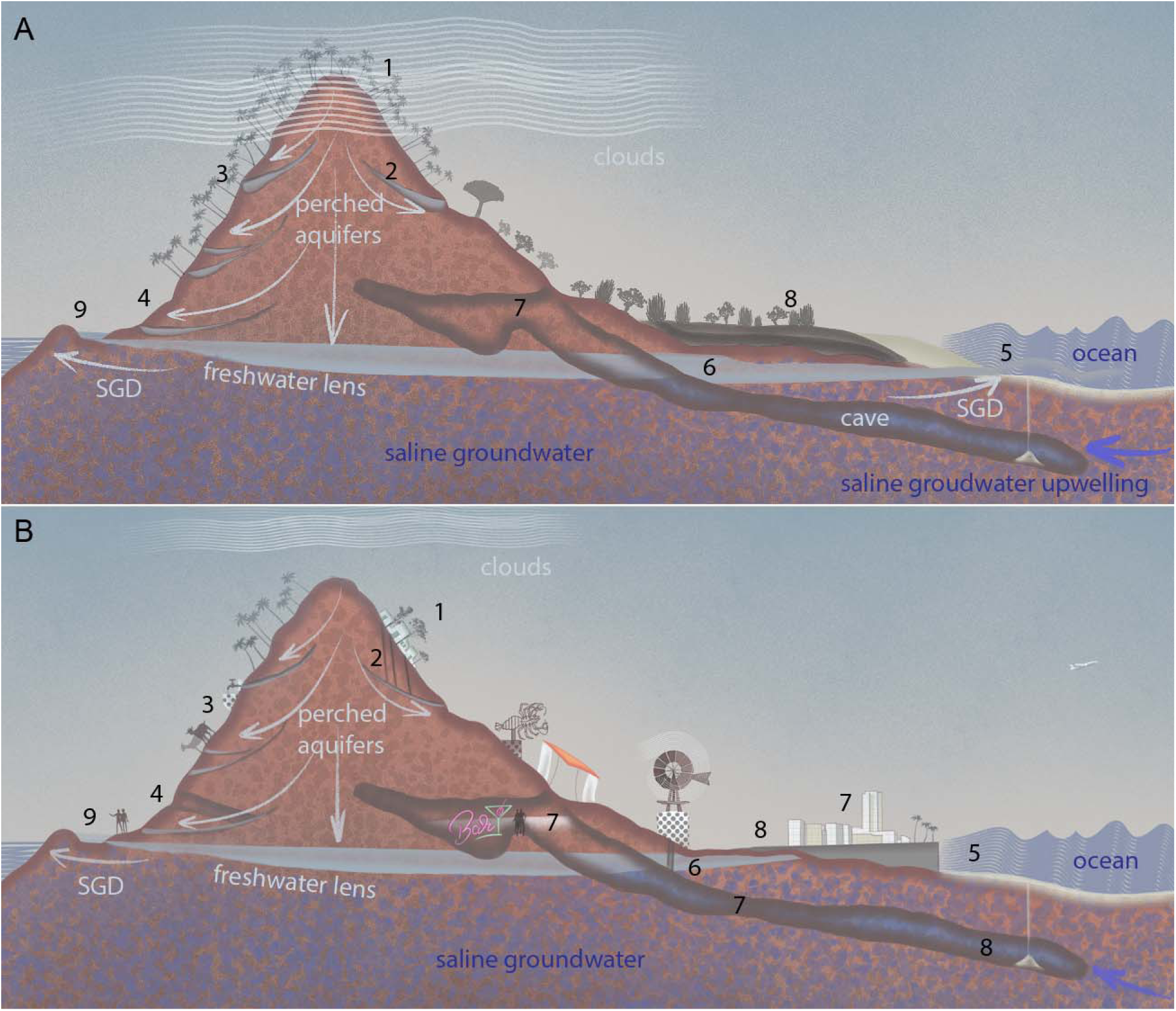
Rationale of the study, summarizing the human-driven impacts on the island aquifer. **A.** Island aquifer in pristine conditions, showing the recharge of the aquifer driven by Tradewind precipitation and hypothetical water flow from highlands to the island’s basal aquifer—partially interrupted by aquitards forming perched aquifers. **B.** Island aquifer in the current state. Urbanization (1) reduces natural thermophilic forest cover and aquifer recharge; (2) wells and cesspits associated with urban areas extract and potentially contaminate perched aquifers; (3) springs associated to aquitards are modified for agriculture and livestock, others (4) are exploited for water mining; (5) coastal development disrupts submarine groundwater discharge, and potentially introduces pollution via illegal wastewater discharge; (6) basal aquifer is exploited via wells for salt production; (7) anchialine caves are altered for tourism; (8) coastal infrastructure degrades vegetation and dune systems, affecting the carbon input into the aquifer; (9) increased population and tourism degrade anchialine pool water quality. Drawing by Guillermo García Gómez.

**Figure 2.**
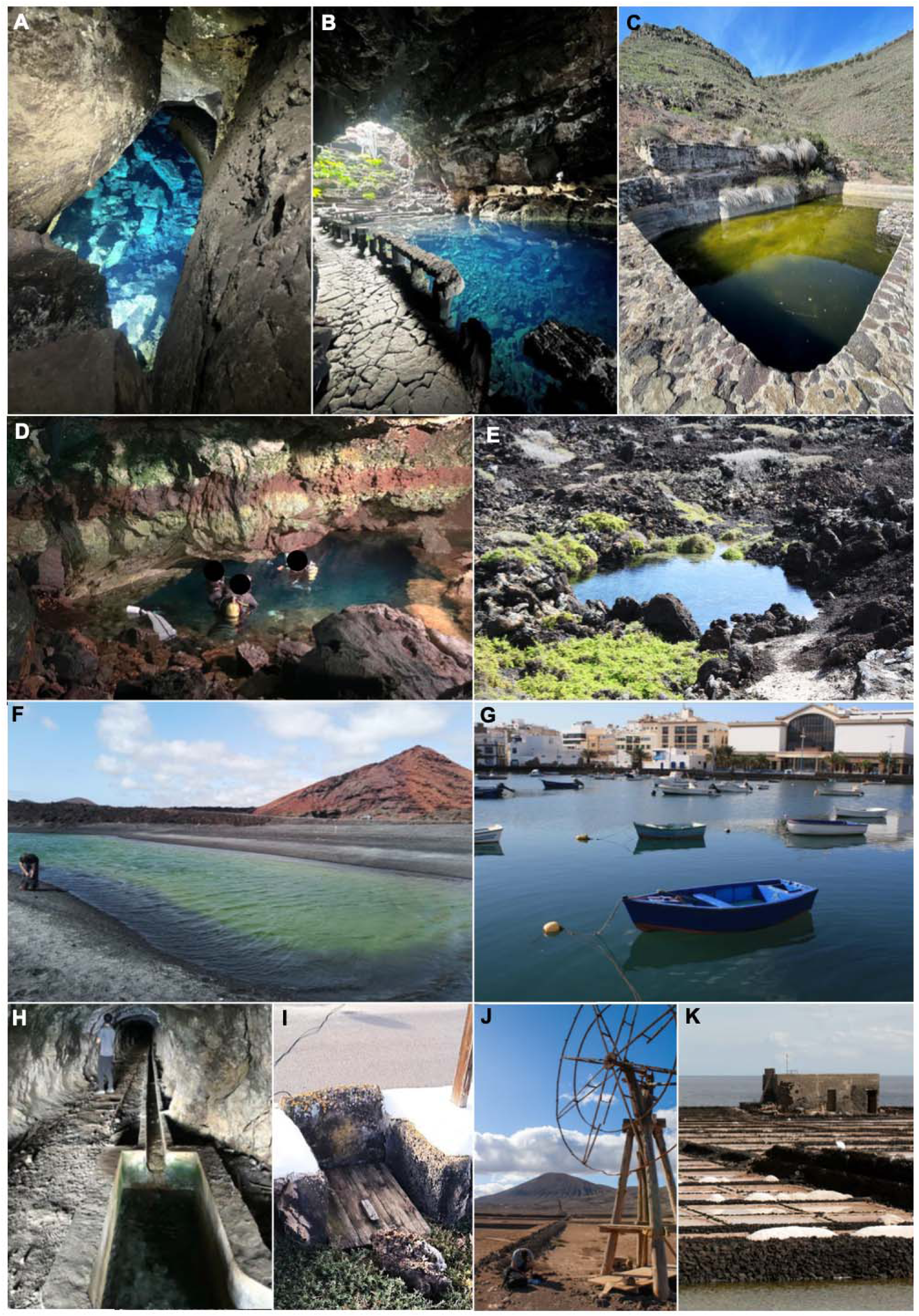
Groundwater and groundwater-dependent ecosystems sampled during this study. **A.** Entrance to the Cueva de los Lagos, a saline, dark, anchialine cave, part of La Corona lava tube. **B.** Los Jameos del Agua, a saline, illuminated anchialine cave, part of Los Jameos del Agua touristic center, within La Corona lava tube. **C.** Fuente del Chafariz, a freshwater spring-fed pool, influenced by rainfall and terrestrial runoff and frequently accessed by livestock. **D.** Entrance to the Túnel de la Atlántida, a saline, artificially illuminated section of La Corona lava tube, part of Los Jameos del Agua touristic center. **E.** Charcos de Luis, a saline, illuminated, anchialine pool in the north of the island, occasionally visited by tourists. **F.** Charco Grande de Montaña Bermeja, a saline, illuminated, anchialine pool in the south of the island, often visited by tourists. **G.** Charco de San Ginés, a modified coastal bay where docks infrastructure has altered submarine groundwater discharge, frequented by fishermen and potentially impacted by pollution by surrounding residential areas. **H.** Famara water mine, accessing the lower part of the brackish perched aquifer of Famara (1-5 ‰). **I.** Urban well in Haría, accessing the brackish perched aquifer of Famara (0.5-0.9 ‰). **J.** Well associated with the saltworks of Los Agujeros, pumping saline groundwater from the island’s basal aquifer into the saltpans for salt production. **K.** Saltpans of Salinas de los Agujeros. People in the images are authors of the paper and they have agreed with publishing their images. Images A-J made by the authors; image K from Wikipedia commons.

**Table 1.** Sampling sites across Lanzarote. The table lists the site name, corresponding sample IDs, habitat classification, and geographic coordinates (WGS 84). Salinity from the samples was measured using a Yinnik multifunction portal sonde; values marked with a tilde (∼) were estimated from sea water; * values measured with refractometers; value highlighted as ** from Wilkens et al, 1993. Notice that in all coastal localities, salinity values vary with the tides (see Wilkens et al. 2009).

| Site | Sample ID | Habitat | Latitude | Longitude | Salinity | Light |
| --- | --- | --- | --- | --- | --- | --- |
| Cueva de los Lagos - entrance | CUL1 | Cave | 29.1577 | -13.4366 | 35.7‰ | Dark |
| Cueva de los Lagos - bottom | CUL2 | Cave | 29.1575 | -13.4343 | 35.7‰ | Dark |
| Los Jameos del Agua | JDA_pool 1<br>(upstream), 2<br>(midstream), 3<br>(downstream) | Cave | 29.1573 | -13.4309 | 35.6‰ | Twilight |
| Túnel de la Atlántida - entrance | TDA_entrada | Cave | 29.1571 | -13.4303 | 36.5‰ | Twilight |
| Túnel de la Atlántida - La Sima | TDA_sima | Cave | 29.1564 | -13.4281 | 36.5‰ | Dark |
| Túnel de la Atlántida - Lago Escondido | TDA_LE | Cave | 29.1565 | -13.4291 | 36.5‰ | Dark |
| Túnel de la Atlántida - Montaña de Arena | TDA_MA | Cave | 29.1575 | -13.4343 | 36.5‰ | Dark |
| Charcos de Luis | CHL | Anchialine pool | 29.2022 | -13.4231 | 40‰** | Light |
| Charco de Punta Prieta | CHP | Anchialine pool | 29.2011 | -13.4211 | 36.5‰* | Light |
| Charco de Suso | CHS | Anchialine pool | 29.1518 | -13.4389 | 38‰ | Light |
| Charco de Zac | CHZ | Anchialine pool | 29.1728 | -13.4266 | 36.5‰* | Light |
| Charco Grande de Montaña Bermeja | MBB | Anchialine pool | 28.9612 | -13.8297 | 38.2‰ | Light |
| Charco Pequeño de Montaña Bermeja | MBS | Anchialine pool | 28.9617 | -13.8266 | 36.1‰ | Light |
| Salinas de los Agujeros - saltpans | CSS 1, 2 | Saltworks | 29.0604 | -13.4614 | 95.8 ‰ | Light |
| Salinas de los Agujeros - pozo | CSP | Saltworks | 29.0608 | -13.4638 | *36‰ | Twilight |
| Caletón Blanco - clean | CBL | Coastal site | 29.2175 | -13.4424 | ~36‰ | Light |
| Caletón Blanco - polluted | CBC 1, 2 | Coastal site | 29.2187 | -13.4444 | ~36‰ | Light |
| Charco de San Ginés | CHG | Coastal site | 28.9612 | -13.5482 | 36.7‰ | Light |
| Punta Usaje | JDA_beach 1 (*littoral*), 2 (*inland*) | Coastal site | 29.1565 | -13.4267 | ~36‰ | Light |
| Casa del Agua de Famara | FWH | Well | 29.1402 | -13.5255 | 2.6 ‰ | Dark |
| Galería de Famara | FWM | Well | 29.1442 | -13.5248 | 2.6 ‰ | Dark |
| Galería del Pescador (Famara) | GAF | Well | 29.1403 | -13.5243 | 3.6 ‰ | Twilight |
| Fuente del Chafaris | FUC | Spring | 29.1231 | -13.5084 | 0.88‰ | Light |
| Pozo de Suso | SHW | Well | 29.1446 | -13.5029 | 0.96 ‰ | Dark |
| Galería de la Fuente del Chafaris | TAB | Well | 29.1231 | -13.5084 | 0.88‰ | Twilight |

**Table 2.** Overview of groundwater-dependent coastal habitats in Lanzarote, summarizing their associated aquifer types, representative sites, main anthropogenic pressures, key ecosystem services, and proposed conservation strategies. For detailed habitat descriptions and additional examples, see Figure 1 and Table 1. (SGD = submarine groundwater discharge; WWTPs = wastewater treatment plants).

| Habitat | Anthropogenic pressures | Ecosystem Services | Proposed conservation actions | References |
| --- | --- | --- | --- | --- |
| *Perched freshwater aquifers (salinity: 0.8 - 3.6 ‰)* |  |  |  |  |
| Mountain and interior wells and spring-fed ponds | Canalization of springs; construction of wells and galleries; groundwater extraction; organic enrichment; pathogens from livestock and human use | Drinking water; wetlands; groundwater dependent vegetation | Monitoring of groundwater extraction; protection of recharge areas; restoration of natural flow paths; sanitation management near springs; regular microbiological and physicochemical monitoring; citizen science monitoring initiatives for private wells; collaboration with Lanzarote's island council | (Custodio, 2002; Izquierdo, 2014) |
| *Basal saline aquifer (salinity: 36.0 - 95.8 ‰)* |  |  |  |  |
| Lowland wells and impermeable pools | Construction of wells, saltworks, and saltpans; water and fauna extraction; artificial wetlands; human presence, pollution, pathogens | Salt production; wetlands (bird breeding) | Regulate groundwater extraction; maintain traditional saltworks sustainably; control pollutants; promote eco-tourism; promote laws for periodical groundwater evaluation; citizen science monitoring initiatives for private wells | (Marín and Luengo, 1994; Masero, 2003) |
| Anchialine caves | Tourist visitation and caving; human pathogens; coin disposal; infrastructure-derived pollution | Tourism; Scientific research; Science communication; Cultural activities | Access control and visitor quotas; regular microbiological and physicochemical monitoring; environmental education; inclusion in protected area management | (Saiz-Jimenez, 2012; Martínez et al., 2020) |
| Anchialine pools | Pollution from visitors, surface runoff, or nearby infrastructure | Temporary fish reservoirs; wetlands; Groundwater dependent vegetation; recreational use | Habitat fencing or visitor control; runoff mitigation; periodic water quality assessment; local stewardship initiatives | (Eshtawi et al., 2016) |
| Submarine groundwater discharge in coastal bays | Nutrient enrichment from SGD; eutrophication risk; contamination (nitrates, pathogens); WWTPs; altered nearshore food webs; degradation of seagrass and rocky reef habitats | Fisheries; coastal productivity; recreation; tourism | Integrated coastal groundwater management; nutrient input reduction; monitoring of SGD plumes; protection of seagrass and reef habitats | (Moore, 2010; Santos et al., 2021) |

Caves were represented by eight samples collected from different sections of La Corona lava tube: Cueva de los Lagos (Figure 2A), Jameos del Agua (Figure 2B), and several sections of Túnel de la Atlántida (Figure 2D, entrance pool, La Sima, Lago Escondido, Montaña de Arena). All provide access to the island’s basal aquifer, which dates back to the island’s formation (15 million years ago), though the lava tube itself formed only 20,000 years ago. These sites span a gradient of human influence, from low visitation at Cueva de los Lagos (50–100 visitors annually) to highly touristic locations in Los Jameos del Agua, inaugurated in 1977 and receiving >800,000 visitors per year (Martínez et al., 2019). Samples from Túnel de la Atlántida were collected along a gradient of increasing distance from the cave entrance at the touristic center, extending to La Sima (∼150 m), Lago Escondido (∼250 m), and Montaña de Arena (∼750 m). Los Jameos del Agua receives limited sunlight whereas, except for its entrance, all cave locations in the Túnel de la Atlántida are in complete darkness (Martinez et al., 2016).

Anchialine pools are tidally influenced inland pools flooded by saline groundwater from the island’s basal aquifer but exposed to light and receiving a strong terrestrial and marine influence through the entrance of particulate organic matter and sea spray. We sampled those at Charcos de Luis (Figure 2E), Charcos Grande and Pequeño de Montaña Bermeja (Figure 2F), Charco de Suso, Charco de Zac, and Charco de Punta Prieta. While most of these pools remain largely undisturbed, Montaña Bermeja is popular for recreational use, and Charco de Zac serves local fishermen as a holding area for bait and fish.

Saltworks were represented by the Salinas de los Agujeros, the only active saltworks still supplied by saline groundwater from the island’s basal aquifer. We collected samples from two hypersaline saltpans (Figure 2K), exposed to sunlight and receiving strong terrestrial input; as well as an abandoned pumping well that showed evidence of fuel contamination (Figure 2J).

Groundwater wells, galleries, and spring-fed ponds were sampled at Tabayesco, Famara, and Haría. These sites access the fresh- or slightly brackish waters of perched aquifers along the mountain ranges of the island, whose formation likely dates back to earlier geological stages, when mountain ridges intercepted moist Tradewinds and hosted dense laurel forests. Today, these wells and mines continue to supply water for domestic and agricultural use, drawing from captured springs, open tanks, and active pumping systems. Wells in Tabayesco and Haría are still regularly used by local farmers and shepherds. The bacterial communities likely reflect a mixture of environmental taxa from both the aquifer and surrounding habitats, as well as anthropogenic taxa introduced through direct human interaction and indirect influences on the aquifer. Local sites in Famara are likely affected by marine spray (Figure 2H), while those in Tabayesco are more influenced by inland conditions and recurrent human and livestock activity. The well in Haría is situated within an urban setting, potentially subject to additional urban-associated impacts (Figure 2I). Although wastewater from Haría is treated at the Haría–Arrieta wastewater treatment plant (WWTP), many houses built before the early 20th century still use independent cesspits instead of connection to the municipal sewer network. Although these habitats differ in environmental context and human impact, all are classified as wells. The only exception was the freshwater spring-fed pond in Tabayesco, which we categorized separately, due to its high exposure to sunlight, rainfall, and external disturbances (Figure 2C).

Coastal environments and enclosed bays influenced by submarine groundwater discharge (SGD) were sampled across three distinct areas of the island, each characterized by a different level of anthropogenic impact. Samples at Caletón Blanco were collected near the small town of Órzola (352 inhabitants, as of 2021), which first appears in maps from the 17th century (Trapero and Santana Martel, 2025) as a small fishermen settlement built around a saltwork complex, nowadays inactive. The village is far from any WWTP and, therefore, the wastewater is likely to be discharged directly into the ocean or collected in fecal wells. At Charco de San Ginés (Figure 2G), located in the island’s capital Arrecife, samples were taken in an enclosed bay within the city. Although the bay is clearly enriched with organic matter from the fishermen’s activities, there is no indication of wastewater discharge in the area, as the city is equipped with functioning WWTP. In contrast, samples at Punta del Usaje were collected within the legally protected area of Malpaís de la Corona, a remote zone with no nearby human settlements and strict regulations prohibiting wastewater discharge.

Water was collected with sterile 5 L containers or sterile ziplock bags during cave dives in the lava tube. All collection materials were pre-sterilized with 5% chlorine and rinsed with sterile distilled water. All samples were processed within 24 hours from collection.

### 2.2. Laboratory procedures and sequencing

Water collected from 30 sampling points was filtered through sterile 0.22 µm pore-size polycarbonate filters (Merck Millipore, Germany) using a peristaltic pump until the filters were clogged. Filters were then stored at −20 °C until DNA extraction. Prior to extraction, each filter was cut into three equal sections, and each section was processed independently to maximize DNA recovery.

DNA was extracted using the DNeasy UltraClean Microbial Kit (QIAGEN, Hilden, Germany), following the manufacturer’s instructions. The concentration of the extracted DNA was measured with a Qubit 1X dsDNA High Sensitivity Assay kit (Thermo Fisher Scientific, Waltham, USA). All DNA samples were subsequently stored at −20 °C until sequencing.

For 16S rRNA gene amplicon sequencing, samples were shipped under controlled conditions to an external service (IGA Technology Services, Udine, Italy). The V3–V4 hypervariable regions of the 16S rRNA gene were amplified using the universal bacterial primers 341F (5’-CCTACGGGNBGCASCAG-3’) and 805R (5’- GACTACNVGGGTATCTAATCC -3’) (Herlemann et al., 2011), and sequenced on a Aviti platform (Element Biosciences, San Diego, CA) using 300-bp paired end mode. Forward and reverse reads were nearly identical in count, as expected from paired-end sequencing.

### 2.3. Sequence processing and dataset assembly

Base calling, demultiplexing, and adapter masking were performed with Element Biosciences Bases2fastq software v.1.8 (https://docs.elembio.io/docs/bases2fastq/). Raw sequences were processed using the DADA2 pipeline (Callahan et al., 2016), in the R environment v 4.5.0 (R Core Team, 2025). Forward and reverse reads were truncated at 280 bp and 220 bp, respectively, after trimming the first 17 and 21 bases to remove primer sequences. Sequences with ambiguous bases were discarded, and the maximum expected errors were set to 2 for forward reads and 4 for reverse reads. Reads with a quality score below 2 were removed, and PhiX contamination was filtered out by setting rm.phix = TRUE in DADA2’s filterAndTrim(), which filters sequences matching the PhiX174 genome. These parameters were chosen based on inspection of quality score profiles using the plotQualityProfile() function. Reads were then merged to generate unique Amplicon Sequence Variants (ASVs). Taxonomic assignment was performed against the Silva database (v138.2, https://www.arb-silva.de/). Raw reads are available in …

We refined the dataset, removing ASVs identified as archaea, mitochondria, chloroplasts, or unassigned at the phylum level (presumed artifacts). We also excluded rare ASVs, defined as those occurring in fewer than three samples or represented by fewer than three reads. The resulting dataset, containing all remaining ASVs regardless of taxonomic assignment, is designated as the ASV table. This dataset was used to describe the taxonomic composition of the bacterial communities across the island, including pathogens, anthropogenic, and environmental bacteria, as well as for the inference on richness and composition patterns across habitats.

### 2.4. Taxonomic composition and richness across Lanzarote groundwater-related ecosystems

In order to describe the pathobiome, as well as the human- and environment-related bacteria, we generated a dataset by clustering ASVs by taxonomy at four ranks—species, genus, family, and order. Out of those clustered operational taxonomic units (OTUs), only those identified to at least the focal rank and represented by ≥50 reads in the dataset were retained. Each taxonomic group was annotated for pathogenic potential based on the nomenclature and taxa related information on pathogenicity levels from the List of Prokaryotic names with Standing in Nomenclature (LPSN; https://lpsn.dsmz.de/) accessed between the 2nd and the 10th of September, 2025 (Table S2). We defined the pathobiome as the total set of genera with potential or confirmed pathogenicity in humans (Pathogenicity level 2 or 3 in the LPSN). Taxa were also classified according to their ecological origin following the predominant habitat information collected from over 4000 sources (metadata related to bacterial isolates, MAGs, and metagenomes) by applying scite_ (https://scite.ai/home) and Copilot AI (https://copilot.microsoft.com/). We classified environment-derived taxa as those for which at least 90% of the reference hits were associated with natural environments. Conversely, we defined human-derived taxa as those for which ≥ 90% of the reference hits corresponded to humans or human constructed environments (including raw sewage). These categories are not mutually exclusive: a bacterial taxon could be pathogenic and have an environmental or anthropogenic origin (Table S2). This classification framework was used to distinguish taxa associated with human-related environments from those adapted to natural habitats, enabling an ecological interpretation of the pathobiome in relation to anthropogenic influence.

### 2.5. Statistical inference

Our overarching hypothesis is that habitats host distinct bacterial assemblages that differ in richness and composition because of two processes: (i) environmental selection associated with contrasting aquifer conditions, which should favour turnover among habitats; (ii) and anthropogenic disturbance of those conditions, which should introduce or remove human-associated lineages and thus increase nestedness.

To distinguish between these two processes, we tested three sub-hypotheses. H1: Taxonomic richness and composition of the whole bacterial community differ among habitats, with compositional differences dominated by turnover (expected under environmental selection); H2: Pathogenic and anthropogenic fractions show habitat-specific enrichment and β-diversity dominated by nestedness (expected under human inputs), whereas the environmental fraction mirrors the whole community and remains turnover-dominated; H3: Differences in species composition are not uniform across habitats. We expect habitats with the most distinct ecological conditions (such as hypersaline saltworks or dark, oligotrophic caves and wells) to host more distinct bacterial communities, thereby increasing their discriminability. In contrast, habitats that are in closer contact with the surrounding environment, such as anchialine pools and coastal sites, should display more overlap. We further expect the discriminability of the entire or environmental community to be higher (reflecting stronger turnover) than that of the anthropogenic and pathogenic bacteria (displaying greater nestedness).

Given strong correlations among environmental and anthropogenic variables (salinity, light, altitude, distance from the sea, number of buildings, and road length), we first assessed collinearity using the function ‘pairs.panels’ included in the R package psych, v. 2.4.1. (Revelle, 2020; |Pearson r| > 0.7). As expected, these variables clustered along two main ecological gradients: an epigean–hypogean axis (surface vs. subterranean habitats) and a sea-land axis (salinity and coastal influence). To reduce redundancy, we retained distance from the sea (km) as a proxy for environmental variation, and the number of buildings within a 1-km radius as a proxy for anthropogenic pressure, while controlling for sequencing depth (reads per sample) as a covariate.

#### 2.5.1 Patterns of taxonomic diversity of the bacterial community across habitats

In order to test our first hypothesis—that the entire bacterial community differs among habitats and that between-habitat dissimilarity is turnover-dominated—we analyzed the richness and composition of the full ASV matrix, without any taxonomical clustering in order to avoid biases linked to the taxonomic coverage of the reference library. We modeled ASV richness with generalized linear models with a negative-binomial family and identity link function using the function ‘glm.nb’ in the R package MASS, v. 7.3 (Venables and Ripley, 2002), which allowed us to account for zero-inflated counting data.

Predictors included habitat type together with continuous covariates representing (i) environmental context: ‘distance from the sea’ in meters, and (ii) anthropogenic context ‘number of buildings within a 1-km radius’. We assessed model assumptions and model fit with the R package performance, v. 0.7.3 (Lüdecke et al., 2021), testing for distribution of residuals, homoscedasticity, multicollinearity and influential observations. Since the model included a set of predictors with both categorical and continuous variables, we summarized the model parameters using type II ANOVA tables produced using the function ‘Anova’ in the R package car v. 3.0.10 (Fox and Weisberg, 2018), which assess the significance of each variable using Wald χ2 tests. We estimated pairwise differences amongst categorical variables by calculating the estimated marginal means followed by a Tukey test, as implemented in the function ‘emmeans’ included in the package emmeans, v. 1.10.7 (Lenth, 2025).

Second, we calculated differences in ASV composition across samples using the function ‘beta’ in the R package BAT v. 2.9.5 using the Bray-Curtis dissimilarity index as well as its nestedness and turnover components (Cardoso et al., 2015). We tested differences in total ASV composition via PERMANOVA, using the function ‘adonis2’ in the R package vegan, v. 2.6-8 (Oksanen et al., 2025) using the same covariate as in the richness model. We visualized the similarity between samples using a non-metric multidimensional scaling ordination (nMDS) using the function ‘metaMDS’ in vegan, as well as hierarchical clustering with average linkage run setting the method "average" in function ‘hclust’ (R Core Team, 2023; R version 4.3.2). We visualized distribution of β-diversity values across samples taken within and across habitat types using density plot, highlighting median values.

#### 2.5.2 Patterns of taxonomic diversity of pathogenic, anthropogenic and environmental bacteria across habitats

We explored whether anthropogenic and pathogenic lineages show habitat-specific enrichment and β-diversity was dominated by nestedness. In contrast, we hypothesized that between-habitat differences in the environmental subset would be mainly driven by turnover.

We modelled the proportion of reads and species corresponding to pathogens, as well as bacteria with anthropogenic and environmental origin using a generalized linear mixed using a framework fitted with function ‘glmmTMB’ included in the package glmmTMB (Brooks et al., 2017), using a negative binomial and the same predictors and diagnostics as above. As above, significance of the model parameters was assessed using Wald χ2 tests via the function ‘Anova’ included in the packager car v. 3.0.10 (Fox and Weisberg 2019) for categorical variables; followed by a Tukey test to assess pairwise-differences calculated with function ‘emmeans’ in the package emmeans, v. 1.10.7 (Lenth, 2022). Those differences were summarized using a heatmap Compositional differences were investigated via PERMANOVA on Bray-Curtis dissimilarities, with NMDS and clustering for visualization. β-partitioning was repeated to compare Total β (Bray-Curtis), turnover, and nestedness components between subsets.

#### 2.5.3 Quantification of discriminant taxa across habitats

We quantified how well bacterial composition discriminates each habitat against the rest using one-vs-rest random forest classifiers (Brieuc et al., 2018). Read counts were aggregated at family, genus, and species ranks. We excluded the spring-fed pond habitat from the random forest analysis since it was represented by only one sample. Models were fit with class-balanced bootstrap sampling and 10,000 trees and evaluated by out-of-bag (OOB) Accuracy and ROC AUC (Area Under the Receiver Operating Characteristic Curve). We repeated the analysis for four bacteria sets: All taxa, Pathogens, Anthropogenic, and Environmental.

## 3. Results

### 3.1. Description of bacterial community composition and richness across Lanzarote groundwater ecosystems

Sequencing produced a total of 2.72 million paired-end reads across the 30 samples. The number of raw reads per sample ranged from 1,226 to 142,323, with an average of 90,748 ± 34,555 reads per sample. We retrieved 2725 ASVs, of which 2241 could be taxonomically assigned to 248 bacterial families and 499 genera, with site-specific dominance patterns (Figure 3,4). The rest (484 ASVs) remained unclassified (Table S1). The families with the highest relative abundance in terms of number of reads across the dataset were Paracoccaceae (12.1%), Flavobacteriaceae (7.8%), Litorivicinaceae (6.1 %), Cryomorphaceae (5.4%), and Pseudoalteromonadaceae (4.7%), followed by Comamonadaceae (4.3%), Cyanobiaceae (4.0%), SAR116 clade (3.9%, order Puniceispirillales) and Rhodothermaceae (2.9%). These families showed clear habitat-specific patterns. For example, Paracoccaceae were common in anchialine pools and coastal habitats, while Comamonadaceae were more abundant in wells.

**Figure 3.**
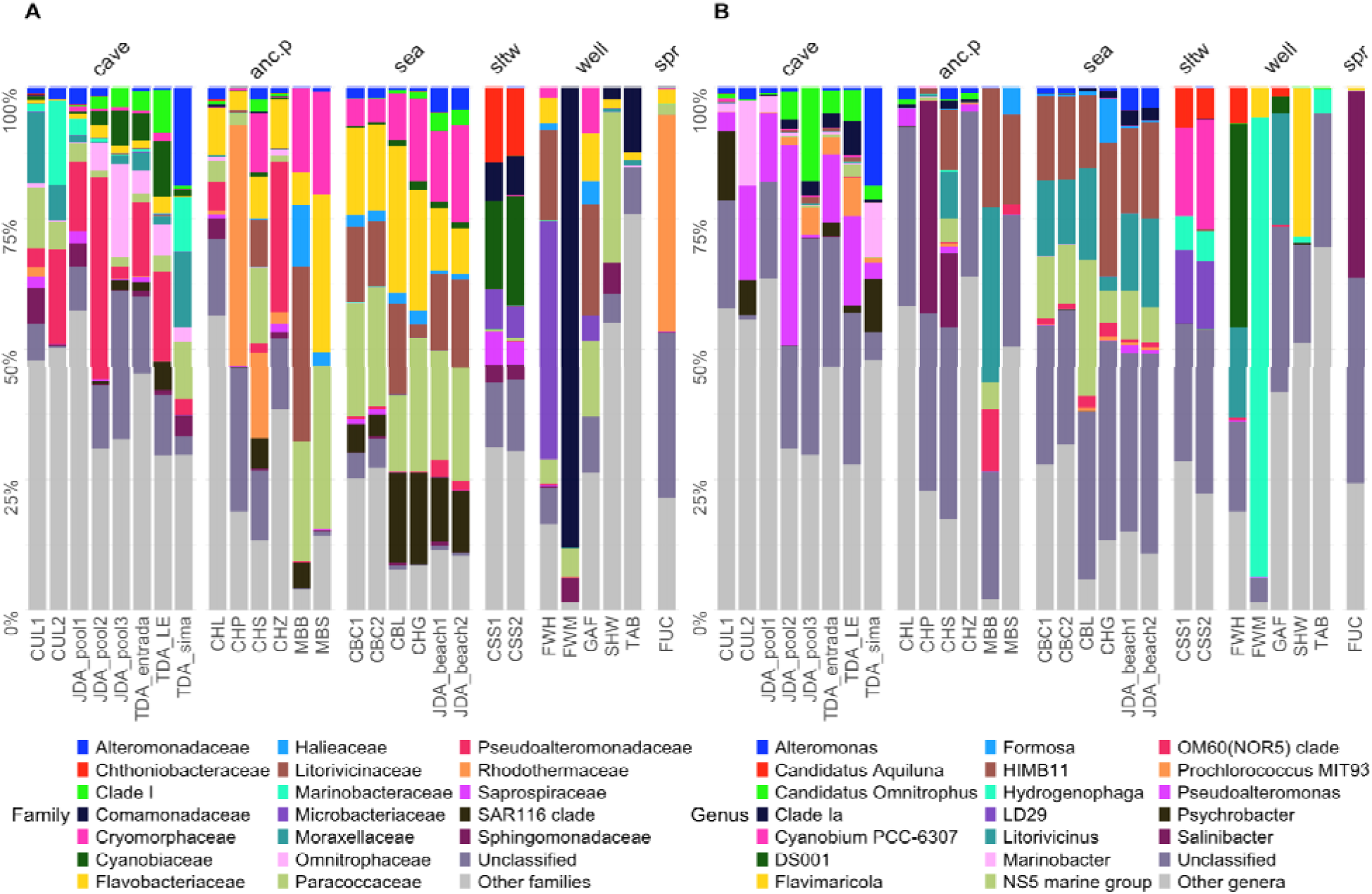
Composition of the entire bacterial community across habitat types, showing relative abundance of the 20 most abundant bacterial taxa across all samples. **A**. Classification at the family rank. **B**. Classification at the genus rank. The category “Other families/other genera” groups those outside the top 20 in abundance, while “Unclassified” indicates sequences not resolved to family or genera rank

**Figure 4.**
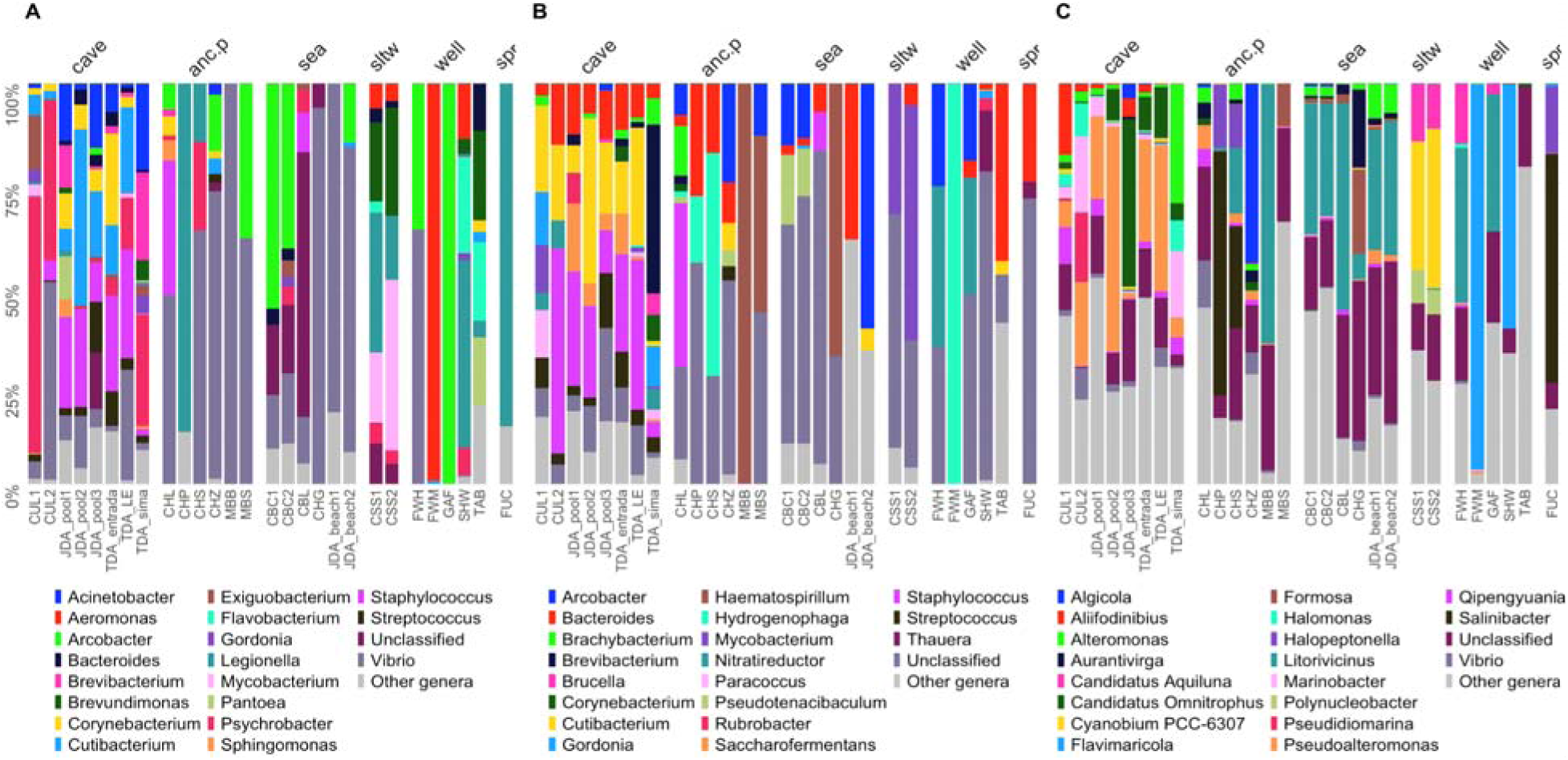
Composition of the pathogenetic, environmental and human-related bacteria across habitat types. **A**. Genera containing OTUs assigned to putative pathogenetic bacteria. **B**. Genera containing OTUs putatively assigned to taxa linked to human or wastewater sources. **C**. Genera containing OTUs putatively assigned to the environment. The category “Other genera” groups those outside the top 20 in abundance, while “Unclassified” indicates sequences not resolved to genus rank.

Cyanobiaceae and Rhodothermaceae were primarily detected in saltworks and in the spring-fed pond, respectively, whereas Pseudoalteromonadaceae were highly represented in cave samples, where richness varied significantly with light and distance from the sea (Tables S6-S7). Unclassified ASVs accounted for 26.4% of the total abundance (Figure 3A).

At the genus level (Figure 3B), the taxa with the highest relative abundances were HIMB11 (Rhodobacteraceae, 7.2%), Litorivicinus (6.5%), Hydrogenophaga (4.0%), the NS5 marine group (Flavobacteriaceae, 5.1%), and Pseudoalteromonas (3.7%). HIMB11 was dominant in anchialine and coastal sites, Hydrogenophaga in wells, and Salinibacter in the spring-fed pond site, while Pseudoalteromonas was the dominant genus in caves. Additional high-abundance genera included Cyanobium (2.8%), Salinibacter (2.4%), and LD29 (Chthoniobacteraceae, 1.9%), each showing strong variation across habitats.

Among families including potential pathogens, Moraxellaceae contributed up to 35.4% of pathogen abundance in cave samples. Vibrionaceae reached 60.5% in anchialine pools and 48.4% in coastal sites. Mycobacteriaceae were more abundant in saltworks, peaking at 33.5% while Legionellaceae represented up to 85.6% of potential pathogen relative abundance in the spring-fed pond sample, 36.1% in wells, and 22.9% in saltworks.

Legionella, Vibrio, Psychrobacter, and Mycobacterium were the most abundant of the 43 identified genera harboring potentially pathogenetic bacteria (Figure 4A). Legionella, in particular, accounted for 85.6% of pathogen relative abundance in the spring-fed pond site and 36.1% in wells; Vibrio accounted for up to 60.5% and 46.0% in anchialine pools and coastal sites, respectively. Psychrobacter was more abundant in caves (27.4%), while Mycobacterium was prevalent in saltworks samples (33.5%). Other clinically relevant genera, such as Enterococcus and Streptococcus, were largely restricted to caves and wells, where they accounted for up to 4.2% of pathogens relative abundance.

Among families with a likely anthropogenic origin, Stapphylococcaceae were the most abundant, reaching up to 20.0% in caves and 18.2% in anchialine pools. Other prominent families included Comamonadaceae (89.9% in wells), Mycobacteriaceae (51.0% in saltworks), and Flavobacteriaceae (14.2% in coastal sites). Bacteroidaceae were the most commonly identified family (24.6%) in the spring-fed pond site. Human-related genera accounted for 5.5% of total bacterial abundance and were dominated by Hydrogenophaga, Staphylococcus, Cutibacterium, Bacteroides, Brevibacterium, and Streptococcus (Figure 4B). These genera were consistently detected in caves, spring-fed ponds, anchialine pools, and wells, where Hydrogenophaga reached up to nearly 90.0% relative abundance. In coastal and saltworks sites, their relative abundance was below 0.5%.

Families to which an environmental origin was assigned were the most abundant group overall, collectively contributing to 50.8% of total bacterial abundance. Pseudoalteromonadaceae were the most common family in cave samples (24.3%), whereas Rhodothermaceae were highly abundant in the spring-fed pond sample (66.4%). Litorivicinaceae reached up to 21.2% in anchialine pools and 29.6% in coastal sites, Cyanobiaceae 36.8% in saltworks, and Paracoccaceae 30.6% in wells. At the genus level, environmental bacteria dominant taxa included Pseudoalteromonas, Litorivicinus, Cyanobium PCC-6307, and Salinibacter (Figure 4C). These genera were very abundant in caves, anchialine pools, saltworks, and the spring-fed pond site, where Salinibacter and Halopeptonella together exceeded 75.0% relative abundance.

Wells and coastal sites were dominated by Flavimaricola and unclassified taxa, at 28.9% and 30.1% of total environmental bacteria relative abundance.

Building on the taxonomic patterns described above, we next assessed how ecological origin and pathogenic potential structure these communities across habitats (Figure 5).

**Figure 5.**
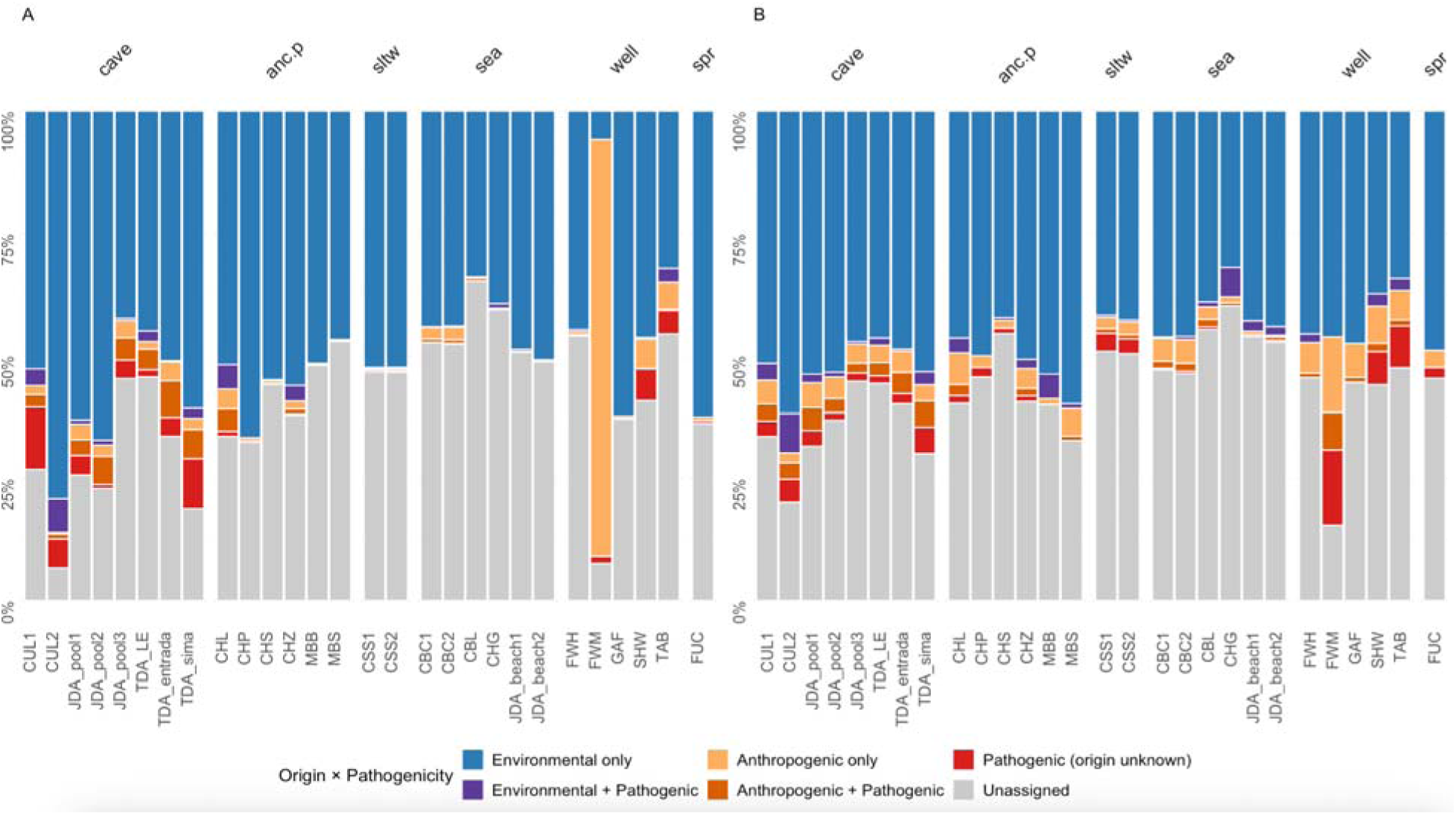
Functional composition of bacterial communities across the samples. **A**. Relative abundance and **B**. taxonomic richness of bacterial taxa categorized by ecological origin (environmental or anthropogenic) and pathogenicity (pathobiome). Each bar represents the aggregate community composition for a specific habitat type.

### 3.2 Inference of habitat-specific patterns in the bacterial community

Bacterial communities differed among habitats in both richness and composition. Taxonomic richness significantly differed across habitats, as confirmed by our generalised linear models (GLM; χ² = 29.702, p < 0.001; Table S3; Figure 6A), regardless of number of reads and other covariates. Pairwise post hoc comparison showed that those differences were only significant between caves and anchialine pools (Tukey test; estimate = 0.816, p = 0.013; Table S4), caves and wells (estimate = 1.594, p <.0001), open sea and wells (estimate = 1.294, p = 0.006).

**Figure 6.**
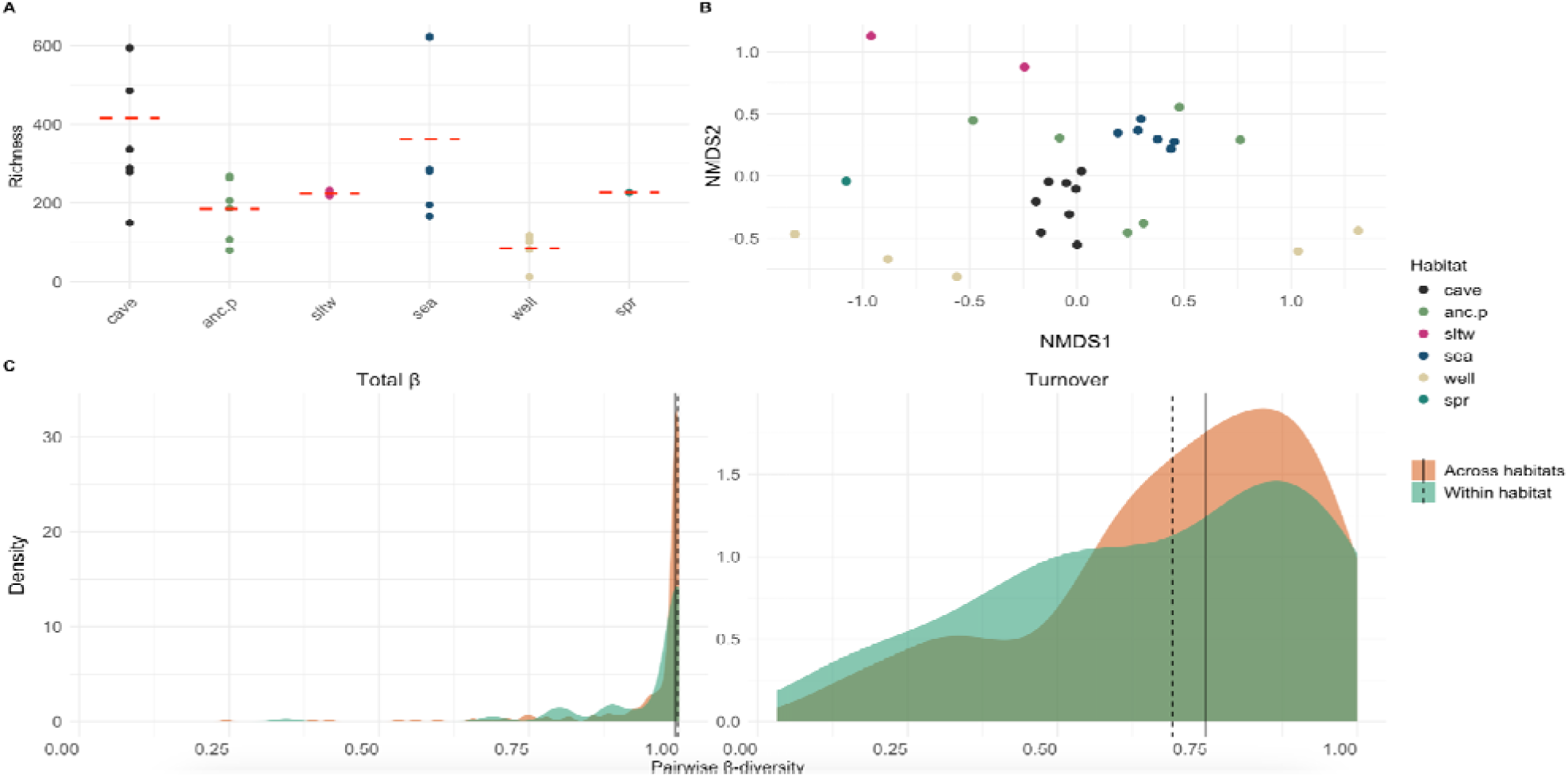
Whole-community bacterial patterns across habitat types. A. Differences in richness across habitats. B. Non-metric multidimensional scaling based on Bray-Curtis distance of whole-community composition; points are colored by habitat type. C. Pairwise β-diversity distributions comparing withinhabitat vs across-habitat pairs, shown for total β and the turnover component; relative nestedness is omitted as it is the complement of turnover (β-nestedness = β-total − β-turnover). (cave = anchialine caves, anc.p = anchialine pools, sltw = saltwork pans, well = wells and water galleries, spr = spring-fed ponds

Total β-diversity amongst samples was high (mean = 0.96), and 22.0% of its variance was explained by habitat type (PERMANOVA, p = 0.022; Table S5). Other covariates did not account for significant variation (all p > 0.05). Partitioning of β-diversity showed that turnover was the dominant component (mean percentage turnover β-diversity = 0.69; mean percentage nestedness β-diversity = 0.31; Figure 6C), although its variation was not significantly explained by any predictor (all p > 0.05) (Table S5). Visualization through NMDS of total β-diversity revealed distinct groups for caves, hypersaline and saltwork habitats (Figure 6B), whereas wells, spring-fed ponds, and anchialine pools overlapped more substantially. Alternative visualization using hierarchical clustering offered comparable results (Figure S3a).

### 3.3 Patterns of taxonomic diversity of bacterial functional groups

Regression models revealed that habitat significantly influenced the proportion of potential pathogen occurrence (GLM, χ² = 104.617, p < 0.0001; Figure 7A, Tables S6-S7), as well as the number of buildings within a 1-km radius (Estimate = 0.177, p = 0.024). In terms of relative richness, potential pathogens were significantly affected by habitat type (GLM, χ² = 59.890, p < 0.0001; Figure 7D; Table S8), along with the number of reads (estimate = 1.059 x 10^-5^, p = 0.039). Community composition analyses indicated clear habitat-specific structuring: PERMANOVA identified significant differences in pathogen assemblages among habitats (R^2^ = 0.221, p = 0.001; Figure 7G, J, Table S10), with nestedness contributing more strongly than turnover (Figure 7G), although none of the variables in our model significantly influenced nestedness (Table S10).

**Figure 7.**
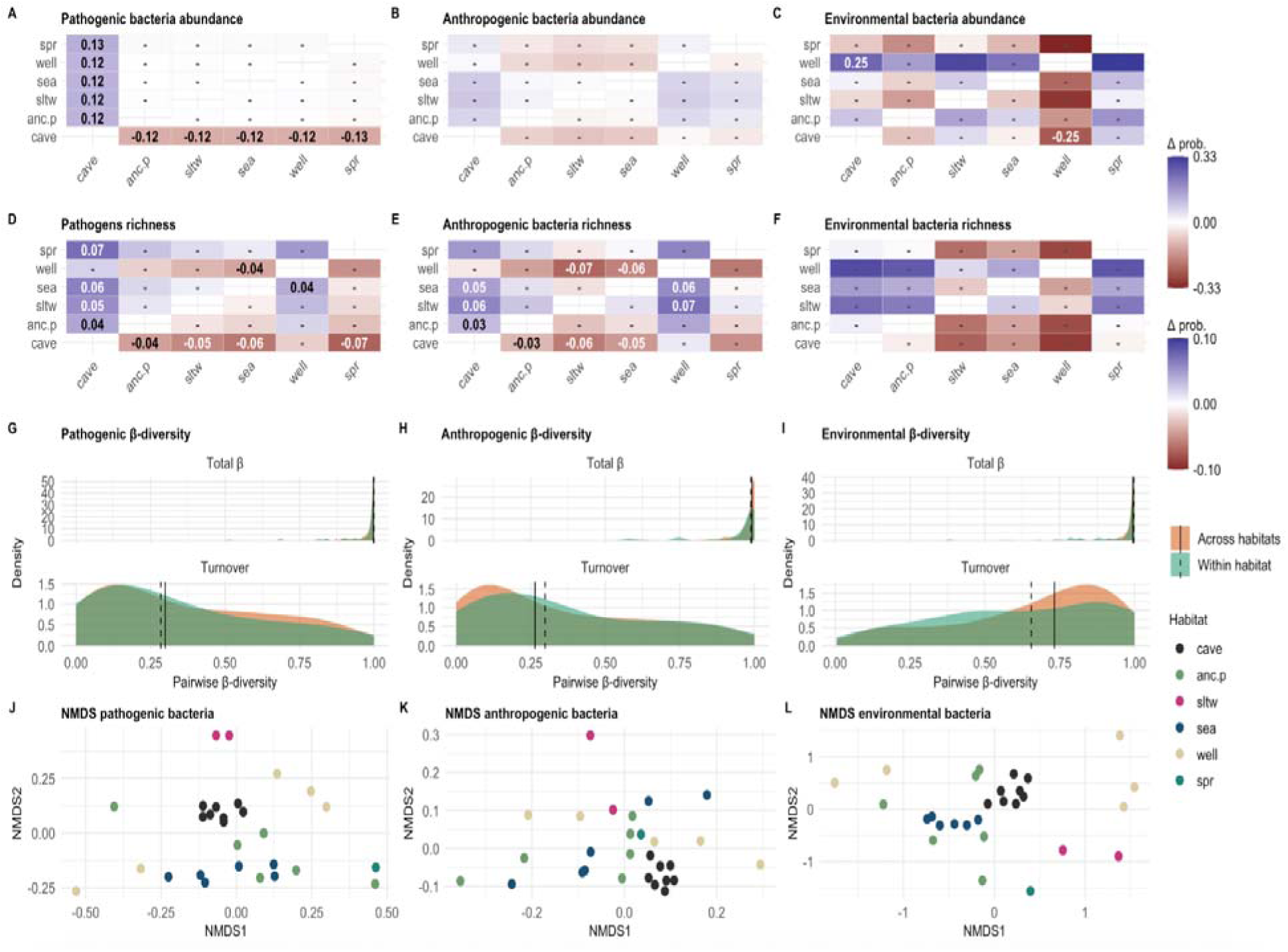
Comparative patterns of bacterial communities across habitat types. A–C Pairwise differences in relative abundance of pathogenic, anthropogenic, and environmental bacteria across habitats. D–F. Pairwise differences in relative richness for the same groups. Blue shades indicate higher values for the row compared to the column; red shades indicate lower values; non-significant contrasts are marked with “–”. G–I. Pairwise β-diversity distributions (β-total and percentage turnover components) comparing within habitat versus across-habitat communities. Relative nestedness is not shown, as it is the specular image of turnover (β-nestness = 1 - β-total). J–L. Non-metric multidimensional scaling (NMDS) ordinations of pathogenic, anthropogenic and environmental bacteria, colored by habitat type. Legends are shared across rows: color scales indicate the range of pairwise differences; shaded density areas show β-diversity comparisons (green = within habitats, orange = across habitats); NMDS colours correspond to habitat types (cave = anchialine caves, anc.p = anchialine pools, sltw = saltwork pans, well = wells and water galleries, spr = spring-fed ponds).

The relative abundance of human-associated bacteria was not significantly affected by any variables included in the model (Figure 7; Table S6-S7). However, their relative richness was significantly influenced by habitat type (GLM, χ² = 33.924, p < 0.0001; Figure 7E, Tables S8–S9). Community composition of human-associated bacteria also differed significantly across habitats (R^2^ = 0.227, p = 0.004; Table S10; Figure 7H, K), primarily driven by nestedness, although, as with pathogens, none of the variables significantly explained nestedness in our models (Table S10).

The relative abundance of environmental bacteria was significantly influenced by habitat type (χ² = 18.495, p = 0.0024; Table S6; Figure 7C), while their relative richness was significantly affected both by habitat (χ² = 15.574, p = 0.0081; Table S8; Figure 7F) and by the number of buildings (Estimate = -0.048, p = 0.0137). Community composition also varied significantly among habitats (PERMANOVA, R² = 0.213, p = 0.040; Table S10; Figure 7L; Figure S3D), with turnover accounting for most of the observed dissimilarity (Table S10).

3.4 Quantification of discriminant taxa across habitats

Overall, saltworks, marine coastal samples, and anchialine caves are well characterized across taxonomic ranks and data subset according to our random-forest analyses. Specifically, saltworks and open coastal habitats showed the most consistent discriminability, followed by anchialine caves (Figure 8). In contrast, the discriminability of anchialine pools and wells was weaker. Rank-wise, discriminability was higher at the family level and followed by genus, whereas classification failed more often at species level. Regarding our community subsets, results based on the whole bacterial community were stronger, followed by those based on OTUs assigned to environmental taxa. Pathogen and anthropogenic models highlighted specific habitats with reduced performance, except for anchialine caves and saltworks.

**Figure 8.**
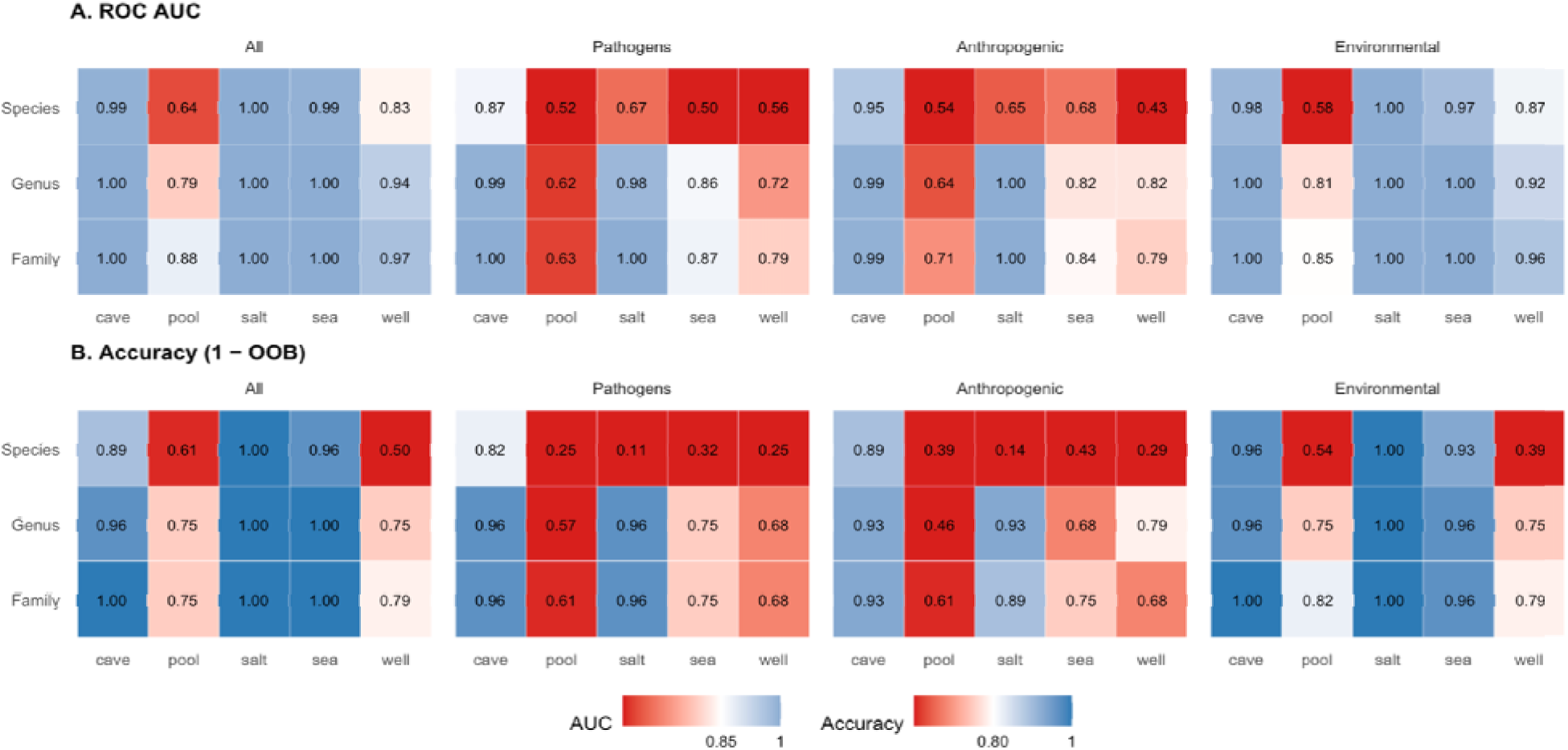
Discriminability of habitats from one-vs-rest random forests across bacterial groups and taxonomic ranks. **A.** ROC AUC. **B.** Accuracy (1−OOB) for models trained separately for each habitat (columns: cave, pool, salt, sea, well) against the remaining habitats, using the entire community (“All”), Pathogens, Anthropogenic, and Environmental subsets. Counts were aggregated to family, genus, and species; spring-fed pond was removed (n = 1). Each tile shows the metric value (10,000 trees; class-balanced sampling). Diverging red–white–blue scales emphasize performance relative to target thresholds (AUC midpoint = 0.85; Accuracy midpoint = 0.80): blue indicates better-than-threshold separability, red worse.

## 4. Discussion

### 4.1 Groundwater ecosystems in Lanzarote are reservoirs of anthropogenic and pathogenetic bacteria

Groundwater-dependent ecosystems in Lanzarote support distinct bacterial assemblages but also exhibit a disproportionate accumulation of human-derived and clinically relevant bacteria (Table S10). Detection of genera linked to fecal contamination raises concerns about the integrity of the island’s aquifer and its capacity to buffer human-derived pollutants. While our sampling design does not pinpoint the exact origins of contamination, the strong association between habitat type and pathogenic enrichment suggests that compartmentalized systems with high water residence time may serve as indicators of anthropogenic pressure.

Cave sites were disproportionately enriched in clinically relevant taxa such as Streptococcus, Psychrobacter, and Mycobacterium, which together accounted for a substantial share of pathogen relative abundance (up to 27 %). We also detected genera linked to fecal contamination, such as Bacteroides and Streptococcus, with the occasional presence of Enterococcus at low relative abundance (Sinton et al., 1993; Teixeira et al., 2020; Li et al., 2021), as well as human-associated genera such as Hydrogenophaga, Staphylococcus and Cutibacterium. These findings support the hypothesis that Lanzarote’s lava tubes, with their partial isolation and limited hydrological renewal, act as sinks of contamination rather than purely natural pathogen reservoirs. This interpretation is consistent with the previous detection of Escherichia coli and Enterococcus spp. associated with the coins thrown by tourists into Los Jameos del Agua cave section, highlighting tourist-linked routes of fecal and anthropogenic bacterial introduction (Martínez et al., 2020). While anthropogenic input appears to be the dominant driver in Lanzarote’s touristic caves, study in a pristine anchialine cave system on the Adriatic coast has shown that resistome diversity can also be structured by natural factors like salinity and stratification, independent of direct human influence (Vojvoda Zeljko et al., 2024).

In contrast, the dominance of environmental specialists such as Pseudoalteromonas, Litorivicinus, and Cyanobium in saltwork and coastal sites, reflects a selective filtering regime likely imposed by chemical determinants like salinity, oxygen, and nutrient availability. This pattern likely emerges from limited anthropogenic enrichment, environmental filtering that limits opportunistic taxa, and hydrological dynamics removing allochthonous bacteria. These factors together might account for the prevalence of specialist lineages in those habitats, in contrast to the anthropogenically influenced communities found in caves and wells.

Pseudoalteromonas species are well known marine specialists associated with surfaces and biofilms, adapted to saline and oligotrophic waters (Holmström and Kjelleberg, 1999; Ivanova et al., 2004), while Litorivicinus and Cyanobium are typical halophilic and phototrophic taxa thriving under high-salinity and high-light conditions in salterns and brines (Oren, 2009). Their presence reinforces the idea that evolutionary lineage sorting operates alongside anthropogenic filtering, generating a dual structuring pattern across Lanzarote’s groundwater ecosystems: one shaped by short-term human disturbance in semi-isolated systems, and another by long-term ecological specialization in open and environmentally filtered habitats.

### 4.2. Turnover dominates whole bacterial assembly, suggesting habitat filtering

Our results demonstrate that bacterial communities in the groundwater-dependent habitats of Lanzarote differ markedly in taxonomic composition, with habitat type emerging as the main driver for turnover (Table S5; Figure S2). This pattern supports the view that bacterial communities in groundwater-dependent ecosystems are primarily shaped by deterministic environmental filtering rather than stochastic or dispersal-limited processes (Stegen et al., 2016; Christensen et al., 2018; Danczak et al., 2018). Similarly, tidal fluctuations and submarine groundwater discharge in a volcanic coastal aquifer might drive predictable, environmentally controlled shifts in coastal microbial assemblages (Lee et al., 2017), whereas salinity gradients in coastal and aquifer systems may act as major environmental filters, structuring distinct bacterial communities across freshwater–seawater transition zones (Héry et al., 2014; Adyasari et al., 2020). These findings support the view that aquifers act as ecological filters shaped by both internal physicochemical heterogeneity and external environmental gradients, promoting high community turnover and low nestedness by selecting for microbial lineages adapted to patchy and often extreme conditions (Kaufmann, 2002; White, 2012).

Analyses of community richness and phylogenetic diversity together revealed that habitat type not only shapes the composition but also the phylogenetic span of bacterial lineages represented within each community (see Supplementary Materials). Caves exhibited significantly higher taxonomic and phylogenetic diversity than wells, consistent with their role as refugia for both diverse and evolutionary distinct clades. In contrast, wells were less diverse, reflecting anthropogenic disturbance, reduced energy supply, and clear evidence of pollution-driven contamination. Other habitats—spring-fed ponds, anchialine pools, and coastal sites—showed intermediate values, suggesting more heterogeneous selective pressures and varying degrees of environmental filtering.

To further resolve these habitat-specific patterns, we partitioned β-diversity into its turnover and nestedness components. At the taxonomic level, between-habitat dissimilarity was dominated by turnover, indicating differentiation through species replacement. From a phylogenetic perspective, however, nestedness played a stronger role, suggesting that although habitats are taxonomically distinct, they are evolutionarily linked as filtered subsets of a deeper, shared phylogenetic lineage (Figure S2; Table S16). This pattern suggests that environmental filtering not only drives taxonomic turnover but also constrains phylogenetic diversity, maintaining evolutionary continuity across habitats. Comparative studies (Bryant et al., 2012; O’Dwyer et al., 2012; Paver et al., 2018) similarly highlight that integrating evolutionary relationships via phylogenetic diversity measurements can reveal filtering patterns not apparent from taxonomy alone, uncovering lineage-specific persistence across environmental gradients.

Comparable phylogenetic structuring has been observed in other subterranean ecosystems, where long-term isolation and extreme physicochemical conditions shape similarly constrained microbial assemblages. In Movile Cave (Romania), long-term chemoautotrophic isolation has produced bacterial assemblages forming a nested subset of global sulfur-oxidizing diversity (Sarbu et al., 1996; Chen et al., 2009). Similarly, Griebler and Lueders (2009) argued that aquifers under low nutrient availability, anoxia, and salinity extremes tend to limit functional and evolutionary diversification, allowing only a restricted set of deep-branching clades to persist. We observe the same pattern in Lanzarote where wells subject to extraction, pollution, and low natural productivity, harbor less distinctive, generalist-dominated assemblages compared to caves, which sustain both higher novelty and greater evolutionary depth.

Our results therefore suggest a dual mechanism. On one hand, taxonomic turnover reflects sharp ecological sorting across habitats, driven by salinity and light regimes (Figure S1). On the other hand, phylogenetic nestedness reflects evolutionary filtering, where only subsets of evolutionarily older (‘deeper-branching’) clades persist under restrictive conditions. Caves, with their unique mix of salinity, organic inputs, and partial isolation, emerge as repositories of both taxonomic novelty and evolutionary diversity. Wells, in contrast, exemplify how anthropogenic modification and nutrient depletion can reduce bacterial assemblages to a smaller set of tolerant, potentially generalist lineages (Tamames et al., 2010).

Ecologically, these findings imply that coastal groundwater-dependent ecosystems function as taxonomic and phylogenetic filters, limiting bacterial diversity not by stochastic loss, but by favoring specific clades adapted to combined stresses of oligotrophy, salinity, and darkness. The contrast between turnover-driven taxonomic divergence and nestedness-driven phylogenetic structuring highlights the need to interpret β-diversity across multiple dimensions: bacterial lineages may be taxonomically distinct, yet they remain evolutionarily constrained, reflecting long-term persistence of specialized clades.

From a conservation perspective, this interplay positions anchialine caves as both refugia of bacterial novelty and repositories of anthropogenic and pathogen-derived taxa. As such, integrating taxonomic and phylogenetic dimensions of bacterial diversity into groundwater management could therefore provide an early-warning framework for contamination, complementing physicochemical monitoring, and supporting the protection of both public health and subterranean ecosystems.

### 4.3. Nestedness dominates human-associated lineages, indicating punctual enrichment

Functional group analyses revealed that anthropogenic and potentially pathogenic bacteria exhibit diversity patterns dominated by nestedness, contrasting with the turnover-driven structure of the overall bacterial community.

Inputs of anthropogenic nutrients and pollutants can substantially alter microbial dynamics in anchialine systems (Kelly et al., 2009; Meland et al., 2023). The interaction of surface runoff with cave waters has been documented to result in microbial transport and deposition within these delicate ecosystems (Davis et al., 2020). These studies, along with our own results, highlight the broader vulnerability of groundwater-fed cave systems to anthropogenic pollution and suggest that such habitats can shift from microbial refugia to pollution sinks under increasing human pressure.

The observation that anthropogenic taxa are enriched in caves but not exclusively confined to them reflects a broader pattern of diffuse bacterial dispersal through coastal aquifers. Rather than being restricted to point-source contamination sites, human-derived bacteria appear capable of spreading across multiple groundwater-fed habitats, likely facilitated by subsurface hydrological connectivity. This is highlighted by the prevalence of potentially pathogenetic and human-related bacteria in remote sections of La Coruna lava tube cave (such as Tunel de la Atlantida and Cueva de los Lagos), which are only occasionally visited by cave divers; and in the water mine of Fuente del Chafariz, which has been closed to visitors for 20 years. Any human signal in those habitats, therefore, most likely reflect infiltration of surrounding soils into the aquifer. Farkas et al. (2022) report that enterococci can survive treatment processes of WWTPs and infiltrate groundwater systems, threatening ecosystem integrity and human health. Their work highlights the inverse relationship between flushing rates and microbial accumulation, suggesting that habitat-specific hydrodynamics, not just proximity to contamination sources, govern microbial load and persistence.

However, it remains unclear whether these potentially pathogenic taxa represent stable cave residents or transient inputs from surface contamination. While some fecal indicator bacteria might survive and replicate in nutrient-poor or dark environments (Felföldi et al., 2016), others might only appear following episodic recharge events, such as rainfall-driven infiltration or increased human presence (Worthington and Smart, 2017; Buckerfield et al., 2019). This ambiguity is echoed in anchialine systems from the Caribbean and Mediterranean, where fluctuating pathobiome signals are often correlated with seasonal or anthropogenic pulses rather than stable community membership (Vucinic et al., 2022).

From a conservation aspect, enrichment of the cave environment with human-derived bacteria is worrisome on two levels: on the one hand it suggests that human impact shifts the bacterial community, introducing markers of anthropogenic pollution with potential impact to human health (Martínez et al., 2020); on the other hand, such shifts might affect higher trophic levels, either through the introduction of allochthonous animal pathogens that could infect the local fauna (Daszak et al., 2001), or through alterations in the food web structure (O’Gorman et al., 2012; Eckert et al., 2019).

### 4.4. The distinctiveness of caves

Cave environments stand out as very distinctive bacterial habitats in our dataset, which regardless of taxonomic rank or functional group, consistently showed high discriminability in random forest models. In our study, cave-associated bacterial communities diverged strongly from those in wells, spring-fed ponds, and anchialine pools (Figures 3,4), underscoring caves as ecological enclaves shaped by unique environmental conditions and limited hydrological connectivity. This is particularly interesting in the case of Tunel de la Atlantida, given that the cave extends along the island’s seafloor and it is subject to strong tidal pumping. Despite this tidal effect, mixing between marine water and saline groundwater remains limited (Wilkens et al., 2009).

Similar degrees of community differentiation have been widely reported in other aquatic cave systems, particularly anchialine caves. For instance, Ghaly et al. (2023) investigated microbial assemblages in Bundera sinkhole, Australia’s only known continental anchialine cave, and found that ∼75% of the metagenome-assembled genomes belonged to novel or uncharacterized families. These communities were stratified along oxygen and salinity gradients, with chemoautotrophic metabolism fueled by nitrogen-sulfur cycling - a clear adaptation to energy-limited, gradient-rich conditions. Similarly, long-term inputs of terrestrial organic carbon have been shown to structure microbial habitats in Bahamian sinkholes.

Sedimentological and geochemical analyses from Blue Hole systems reveal that centuries of terrestrially derived organic matter deposition shape redox stratification and resource availability in these environments (Kovacs et al., 2013; Tamalavage et al., 2018).

Complementary 16S surveys in Bahamian anchialine caves further demonstrate strong microbial stratification along salinity and oxygen gradients (Gonzalez et al., 2011). Other studies have emphasized the extreme heterogeneity of anchialine bacterial communities, even across geographically close sites. For instance, Havird et al. (2022) reported highly distinct microbial communities among nearby Hawaiian anchialine habitats such as Hanamanioa (Maui), Skippy’s Pond (Maui), and Pohue Bay (Hawai‘i Island). Similarly, Kajan et al. (2022) observed that microeukaryotic and prokaryotic communities differed significantly across four anchialine caves—Vjetruša, Blitvica, Gravrnjača, and Živa Voda—in Kornati National Park, Eastern Adriatic Sea, despite their close geographic proximity. Mejía-Ortíz et al. (2022) highlighted the sensitivity to small-scale fluctuations in salinity, oxygen, and organic matter, of bacterial assemblages in anchialine systems, responding sharply to subtle perturbations despite overall habitat stability.

These characteristics mirror our findings in Lanzarote caves, where bacterial communities differ markedly from those in other habitats, exhibiting high taxonomic turnover. The dominance of turnover in shaping community differences is consistent with patterns observed in anchialine systems of the Yucatán Peninsula, where bacterial assemblages are strongly filtered by salinity and nutrient gradients (Pohlman, 2011; Brankovits et al., 2017). However, Lanzarote caves also accumulate human-derived bacteria, pointing to aquifer contamination likely driven by tourism infrastructure and historical groundwater exploitation (Izquierdo, 2014). This contrast suggests that while the ecological drivers of bacterial diversity may be universal, the anthropogenic overlay differs among volcanic islands. Thus, our results echo broader patterns in subterranean microbiology and reinforce the ecological value of cave habitats as reservoirs of bacterial diversity.

Other distinctive habitats in our study were—unsurprisingly—saltworks, where extremely high salinity (∼95‰) probably selects for highly specialized halophilic taxa and excludes both generalist and anthropogenic bacteria (García Jiménez et al., 2020). These systems had high turnover, low nestedness, and high proportions of environmental taxa. They also showed strong separability in random forest models, particularly at the family and genus levels, supporting our hypothesis’ prediction that strong environmental filtering enhances community distinctiveness.

By contrast, anchialine pools were among the least distinct. These ecotonal habitats, exposed to both terrestrial runoff and marine influence, showed greater taxonomic overlap with other systems and lower classification accuracy. Their β-diversity profiles also revealed moderate nestedness and reduced turnover, while their relative richness and abundance of both environmental and anthropogenic taxa were intermediate. This is consistent with our expectation that transitional environments—due to their openness, environmental mixing, and intermediate conditions—would exhibit more diffuse filtering and reduced microbial separation.

### 4.5. Towards effective protection of coastal aquifers considering hydrological and biological components

The groundwater-dependent habitats of Lanzarote—including wells, spring-fed ponds, anchialine caves and pools, saltworks, and coastal discharge zones—support a diverse array of ecosystem services, ranging from freshwater provision and salt production to fisheries, coastal productivity, and cultural value. However, this multifunctionality is paired with a pronounced ecological fragility. Each habitat functions as a finely balanced system of geological, hydrological, and biological interactions, which can be readily disrupted by contamination or hydrological change. Even minimal anthropogenic inputs may trigger cascading effects across microbial and faunal networks, ultimately degrading both ecosystem integrity and water quality (Table 2).

The detection of human-derived and clinically relevant bacteria (e.g., Enterococcus, Streptococcus) highlights that aquifer-dependent ecosystems not only sustain bacterial diversity but can also act as reservoirs of contamination and potential public health risks. Together, these findings underscore the dual ecological role of island aquifers: as reservoirs of bacterial diversity and as sinks of human-derived micro-pollutants. From a management perspective, caves and wells should be treated as sentinel habitats for monitoring human pressures on groundwater. Integrating bacterial diversity and pathobiome assessments into conventional water-quality programs could provide early-warning indicators of aquifer degradation, helping to safeguard both subterranean biodiversity and public health in vulnerable island systems.

Future studies should aim to incorporate seasonal sampling to capture potential temporal shifts in bacterial diversity and community structure, offering a more comprehensive baseline for long-term monitoring.

On the European scale, the Water Framework Directive (WFD) and its related instruments provide the legal foundation for groundwater management, but biological criteria remain largely absent from their implementation (European Commission, 2000). Recent discussions on the revision of the WFD and Groundwater Directive have again drawn attention to this gap (Mammola et al., 2019; Di Lorenzo et al., 2024). Despite those actions, biological indicators are still not formally required, leaving most subterranean ecosystems outside the scope of environmental status assessments. In the Canary Islands, groundwater governance follows the Spanish national Ley de Aguas (RDL 1/2001) and the regional Ley de Aguas de Canarias (12/1990), which delegate management to the Consejos Insulares de Aguas. These authorities prepare and enforce Insular Hydrological Plans that define quantitative and chemical objectives for each island’s aquifers. However, the plans focused on water chemistry and rarely address the biological dimension of groundwater or its connections with coastal ecosystems. To further complicate the situation, subterranean aquatic habitats, such as anchialine systems (caves and pools), perched aquifers, and submarine groundwater discharge zones are only partially mapped, despite their ecological and socio-economic importance (Oromí et al., 2021).

Moreover, monitoring networks are unevenly distributed, and routine assessments of microbiological quality or nutrient fluxes to coastal waters are still lacking. These deficiencies can be corrected within the existing legal framework by expanding monitoring and management toward an explicit bio-hydrological approach. Key improvements for Lanzarote and other islands might include (Table 2):

1. Incorporating microbial and faunal indicators (e.g. diversity indices, pathogen screening) in groundwater-status assessments.
2. Mapping and protecting subterranean aquatic habitats in the Insular Hydrological Plan, with provisions for visitor control and recharge-zone protection. Extending nutrient and microbiological monitoring to submarine groundwater-discharge plumes, where eutrophication and contamination reach coastal reefs and seagrass beds.
3. Aligning groundwater management with protected-area planning, ensuring that wells, ponds, and caves inside or near Natura 2000 or UNESCO Geopark sites receive coordinated protection.
4. Creating a public inventory of groundwater-related wetlands and anchialine sites, following Ramsar principles and open-data standards.

By coupling hydrological, chemical, and biological monitoring, Lanzarote could serve as a model for ecosystem-based groundwater governance in outermost EU regions. Such integration is essential to safeguard subterranean biodiversity, maintain ecosystem services, and ensure the long-term sustainability of the island’s groundwater and groundwater-related resources.

## Acknowledgments

We thank the Cabildo de Lanzarote and the Centros de Arte, Cultura y Turismo (CACT) for their 758 support. We are especially grateful to the staff at Jameos del Agua, particularly Suso Fontes and 759 Alexis Guadalupe, for facilitating access and fieldwork at the site. We also thank GISMACAN S.L., 760 especially Enrique Domínguez, for their support and administrative coordination of the project.

## Data availability

The sequencing data have been deposited in NCBI and are publicly available. The project 793 number is provided in the manuscript. Code is available at https://github.com/amartinezgarcia/DiNezioetal_Lanzarote16S.

