## Supplementary Materials for "Multilayered human activities shape the microbial communities of groundwater-dependent ecosystems in an arid oceanic island"

^11^ Lanzarote and Chinijo Islands UNESCO Global Geopark, Monumento al Campesino, Patio de Artesanos, Ctra. Arrecife a Tinajo 8, 35559, Mozaga, Lanzarote, Spain

^12^ Global Change Research Institute (IICG), Rey Juan Carlos University, C/ Tulipán s/n, 28933, Madrid, Spain

**Supplementary Text 1**

**Microbial dynamics in cave ecosystems**

To investigate habitat-specific dynamics in greater depth, we conducted a focused analysis of cave microbiomes, treating these semi-enclosed environments as ecological enclaves. Caves displayed distinct patterns in microbial richness, composition, and functional group structure, influenced primarily by light availability and proximity to the sea. We followed the same methods as for the entire dataset, but including the following covariates:

A likelihood ratio test (Table S11) revealed that both light regime (p < 0.001) and distance from the sea (p = 0.005) significantly predicted bacterial richness. Pairwise comparisons (Table S12) showed that twilight zones - areas with partial light exposure - hosted significantly more diverse communities than either fully dark (p = 0.0002) or fully sunlit sites (p < 0.0001). These intermediate-light areas appear to act as microbial diversity hotspots, likely benefiting from stable humidity, moderate temperatures, and low disturbance. Richness also declined with increasing distance from the sea, suggesting that marine connectivity may contribute to cave microbiome structure, potentially through saltwater intrusion or coastal runoff.

Despite these richness gradients, community composition remained relatively stable along cave transects. PERMANOVA revealed that none of the measured environmental variables significantly influenced total β-diversity (all p > 0.1; Table S13), indicating spatial consistency in microbial assemblages. However, partitioned β-diversity analyses showed a marginal effect of light on nestedness (p = 0.064), suggesting that light availability may influence selective loss or retention of taxa, rather than community turnover per se.

Importantly, caves harbored the highest load of pathogenic and human-associated bacteria across all habitats. Several clinically relevant genera were frequently detected in multiple cave subsites, including both entrances and deeper twilight areas. Pathogen prevalence was markedly higher in caves than in open aquatic systems, including coastal and saltworks habitats, which consistently showed low or negligible pathogen signals. Hierarchical clustering based on pathogenic and anthropogenic taxa (Figure S3) further emphasized this pattern, consistently grouping cave samples apart from cleaner marine environments.

**Supplementary Text 2**

**Phylogenetic diversity calculation**

To estimate phylogenetic diversity among bacterial assemblages, we reconstructed a phylogenetic tree from the aligned 16S ASVs sequences using the MAFFT version 7 (Katoh et al., 2019), allowing the software to choose the optimal alignment optimal strategy based on our dataset. We then compute pairwise genetic distances and a neighbor-joining tree using the functions dist.dna and njs from the package ape version 5.0 (Paradis and Schliep, 2019). We set the option pairwise deletion as true to handle missing data. Faith’s Phylogenetic Diversity was then computed for each sample using the function ‘alpha’ included in BAT version 2.9.5 (Cardoso et al., 2015). To assess patterns of phylogenetic composition among samples, we calculate pairwise β-diversity metrics using the function ‘beta’ in BAT. To evaluate environmental correlations of phylogenetic richness, we implemented generalized linear models with a Gamma distribution and log link, using the same model structure as for the taxonomic diversity (see Methods). Model assumptions were checked using ‘check_model’ function in the package performance version 0.11 (Lüdecke et al., 2021). Pairwise post hoc contrasts between habitat types were computed with Tukey adjustment using the emmeans package. Differences in total, nestedness, and turnover ASVs composition components across habitat types were tested using permutational multivariate analysis of variance (PERMANOVA) via using the function adonis2 in vegan (Oksanen et al., 2025) using the same model as above. We set 9999 permutations and “by = terms”. We visualize within- and between-habitat differences in β-diversity using density plots.

**Phylogenetic diversity results**

Phylogenetic diversity (PD) also varied significantly with habitat type (χ² = 14.7, p = 0.012; Table S14) with all the other covariates having no significant effect (all p > 0.2). Post hoc pairwise comparisons showed that caves harbored higher PD than wells, reflecting both deeper evolutionary lineages and greater overall richness. All other comparisons were non-significant (Table S15). Phylogenetic β-diversity further supported the structuring role of habitat. PERMANOVA revealed a significant effect of habitat type on total phylogenetic β-diversity (p = 0.011; Table S16), indicating that microbial lineages vary substantially among environments.

Unlike taxonomic β-diversity, which was driven by turnover, phylogenetic differences were more influenced by nestedness, although marginally non-significant (p = 0.063). This suggests that microbial communities across habitats remain evolutionarily connected, but selective filtering leads to the persistence of only specific clades under restrictive conditions. Wells retained only shallow subsets of lineages, while caves preserved phylogenetically diverse assemblages.

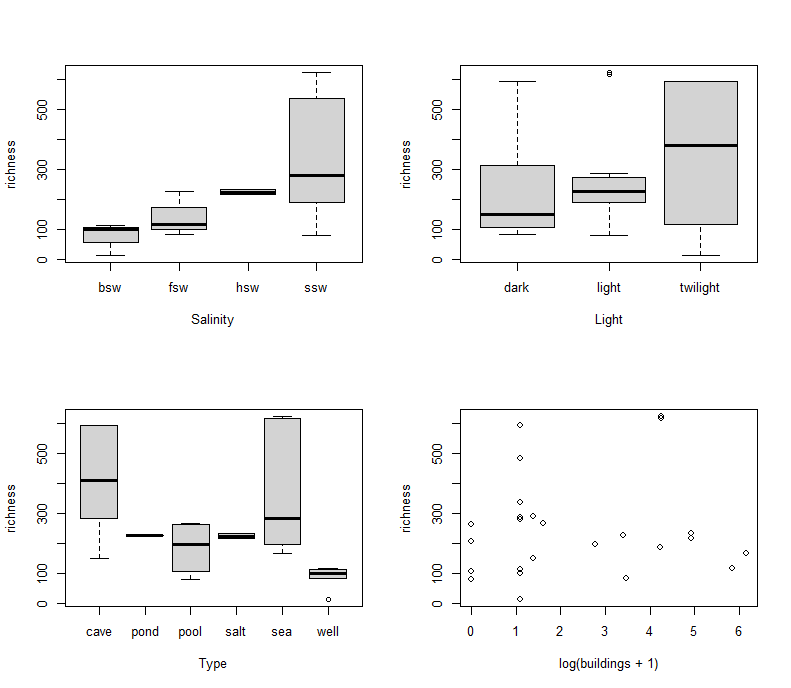

**Figure S1. Patterns of bacterial richness across environmental and anthropogenic gradients. (a)** Richness values across salinity categories: brackish seawater (bsw), fresh seawater (fsw), hypersaline seawater (hsw), and saline seawater (ssw); **(b)** Effect of light exposure on richness; **(c)** Richness across habitat types; **(d)** Relationship between richness and urbanization, shown as log-transformed number of buildings within 1 km.

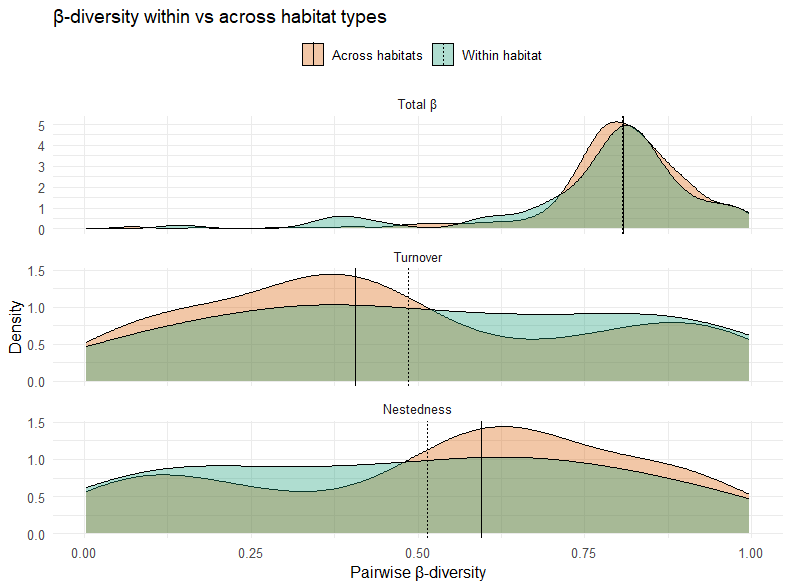

**Figure S2.** **Partitioning of phylogenetic β-diversity within and across habitat types.** Density plots of pairwise β-diversity values for total β (top), turnover (middle), and nestedness (bottom) components. Distributions compare comparisons within the same habitat type (cyan) vs. across different habitats (orange).

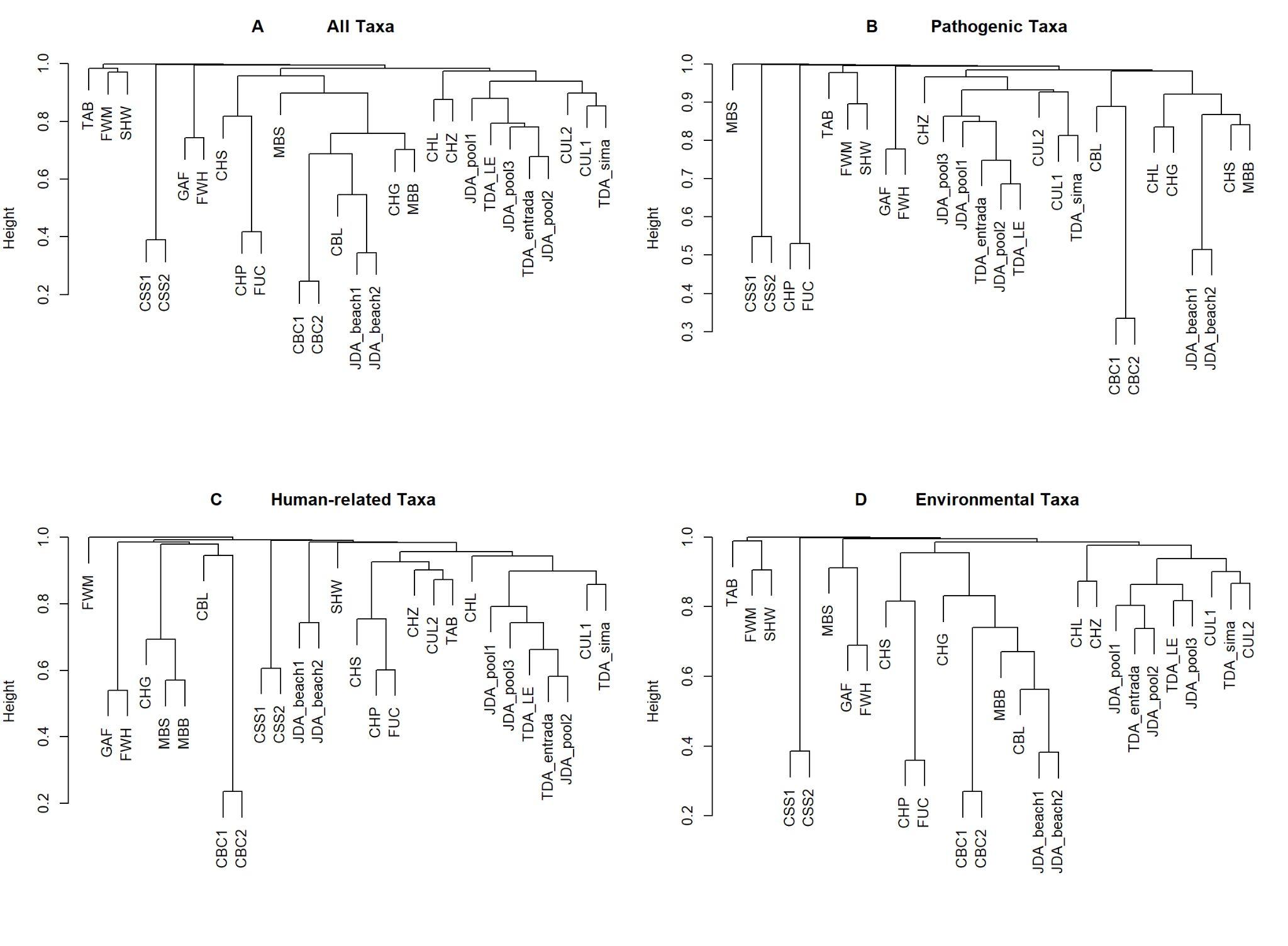

**Figure S3.** Dendrograms show similarity in composition across samples for: **A** the whole bacterial community, **B** pathobiome, **C** human-related and **D** environmental bacterial taxa.
